# Vernalization alters sugar beet (*Beta vulgaris*) sink and source identities and reverses phloem translocation from taproots to shoots

**DOI:** 10.1101/2020.01.28.922906

**Authors:** Cristina Martins Rodrigues, Christina Müdsam, Isabel Keller, Wolfgang Zierer, Olaf Czarnecki, José María Corral, Frank Reinhardt, Petra Nieberl, Frederik Sommer, Michael Schroda, Timo Mühlhaus, Karsten Harms, Ulf-Ingo Flügge, Uwe Sonnewald, Wolfgang Koch, Frank Ludewig, H. Ekkehard Neuhaus, Benjamin Pommerrenig

## Abstract

During vegetative growth, biennial sugar beets maintain a steep gradient between the shoot (source) and the sucrose-storing taproot (sink). To shift from vegetative to generative growth, they require a chilling phase, called vernalization. Here, we studied sugar beet sink-source dynamics upon cold temperature-induced vernalization and revealed a pre-flowering taproot sink to source reversal. This transition is induced by transcriptomic and functional reprogramming of sugar beet tissue, resulting in a reversal of flux direction in long distance transport system, the phloem. As a key process for this transition, vacuolar sucrose importers and exporters, BvTST2;1 and BvSUT4, are oppositely regulated, leading to re-mobilization of sugars from taproot storage vacuoles. Concomitant changes in the expression of floral regulator genes suggest that the now deciphered processes are a prerequisite for bolting. Our data may thus serve dissecting metabolic and developmental triggers for bolting, which are potential targets for genome editing or breeding approaches.

## Introduction

Plants modulate not only the shape and size of their organs, but also physiological and molecular properties in these structures during development and as a response to environmental stimuli. In general, sink organs in plants depend on the import of carbohydrates, mainly sucrose, from source organs. However, previous sink organs may differentiate into ‘sources’, which then, in turn, provide mobilized storage products to newly emerging sinks.

The relative strengths of sinks and sources can be adjusted by the activity of sucrose synthesizing and degrading enzymes (Herbers and Sonnewald, 1998), and by alteration of the activities of phloem located sucrose loaders (Imlau et al., 1999; Gottwald et al., 2000; Srivastava et al., 2008; Chen et al., 2012). As a consequence, both, sucrose metabolizing enzymes and transporters represent targets relevant for breeding strategies aiming at yield increase of crops (Ludewig and Sonnewald, 2016; Sonnewald and Fernie, 2018).

Sucrose is the primary sugar transported in the phloem from source to sink organs. After unloading at the sinks, sucrose can be used as energy precursor, and as building block for growth and storage compound biosynthesis. Non-green storage organs like tubers or taproots must maintain a steep source to sink gradient. To do so, imported sucrose is rapidly converted into relatively inert storage compounds like starch or is compartmentalized intracellularly into large cell vacuoles. As given, sink and source identities of plant organs are dynamic and corresponding transitions are initiated after onset of endogenous developmental signals (Turgeon, 1989) or in response to specific environmental stimuli (Roitsch, 1999). Thus, dynamic regulation of genes and enzymes involved in carbohydrate metabolism and import of sugars into the phloem of mobilizing storage organs are key for source establishment of former sinks (Viola et al., 2007; Liu et al., 2015; O’Neill et al., 2013; Boussiengui-Boussiengui et al., 2016).

Sugar beet (*Beta vulgaris*), the major crop species providing industrial sucrose in the temperate zones of Europe and North America, exhibits a biennial lifecycle and forms a large taproot during the first year of its development. This taproot represents a reversible sink, which contains up to 20% of its fresh weight as sucrose. The vacuolar sucrose loader, named <u>T</u>ONOPLAST <u>S</u>UGAR <u>T</u>RANSPORTER2;1 (*Bv*TST2;1), has been identified to be a key element for sugar accumulation in this storage organ (Jung et al., 2015). During the second year the taproot provides previously stored sucrose as precursor for the formation of a markedly large inflorescence.

The emergence of the sugar beet inflorescence strictly depends on a previous phase of cold temperatures, which induces molecular reprogramming known as vernalization. This vernalization-dependent bolting leads to significant loss of taproot sugar and biomass, and therefore yield. This loss of yield contributes to the fact that sugar beets are solely cultivated as an annual crop. Accordingly, sugar beet is sown in spring and harvested in the following late autumn. A prolonged cultivation period (particularly autumn to autumn) and thus, identification of bolting-resistant varieties have therefore become primary goals in sugar beet breeding over the last decades (Hoffmann and Kluge-Severin, 2011; Hoffmann and Kenter, 2018).

Two major early-bolting loci, *B* and *B2* have been identified in the sugar beet genome in recent years, encoding the pseudo response regulator gene *BOLTING TIME CONTROL 1*, *BTC1* (Pin et al., 2012) and the *DOUBLE B-BOX TYPE ZINC FINGER* protein *BvBBX19* (Dally et al., 2014), respectively. In annual beets, expression of both genes leads to repression of the floral repressor gene *FT1*, and subsequent induction of the floral inducer gene *FT2* and vernalization-independent flowering upon long-days (Pin et al., 2010; Dally et al., 2014). Biennial beets are homozygous for the recessive *btc1* and *bbx19* alleles, which encode non-functional proteins unable to repress the inhibitory function of FT1 (Pfeiffer et al., 2014). Accordingly, biennial sugar beets require vernalization for BTC1- and BBX19-independent *FT1* repression and flowering (Pin et al., 2010). Obviously, floral induction and sink-source transition must be tightly interconnected in sugar beet. A coordinated network of floral inducers and repressors initiates the transition to bolting after vernalization, but adjustment of the metabolic set-up appears equally important for the morphological and physiological restructuring of taproots prior to formation of inflorescences. However, little information is available on the molecular physiological processes in sugar beet at the early time points of vernalization.

In this work, we therefore sought to understand how chilling temperatures, representing a *condition sine qua non* for vernalization, might influence sugar metabolism, photosynthesis, phloem translocation, and therefore source and sink identities of shoots and taproots. We combined comprehensive transcriptome and proteome analyses with recording of organ growth characteristics, photosynthetic parameters and metabolite quantification. In summary, our analyses revealed an unexpected cold-dependent reversal of sink and source identities of taproots and shoots, respectively, prior to bolting at the very early stages of vernalization.

Despite inactivation of photosynthesis in the cold, shoot biomass increased at the expense of taproot sucrose. We recorded a substantial export of taproot sugar in the cold, which correlates with altered activities of sugar ex- and importers and with a markedly altered expression of genes involved in either sucrose synthesis or degradation. We speculate that this so far hidden metabolic reprogramming is a prerequisite for initiation of bolting as corresponding flux redirection transports sugars from the taproot to the shoot. However, this process might also contribute to the pronounced frost sensitivity of sugar beet. Thus, our findings provide a molecular-physiological explanation to the well-known problem of sugar beet cultivation (loss of yield due to the biennial lifecycle) and provide new targets to achieve bolting resistance and winter hardiness in this crop species.

## Results

### Cold exposure causes rapid loss of shoot and root water, but not of shoot biomass production

To resolve cold-dependent growth dynamics of sugar beet source and sink organs, we monitored shoot and taproot weights of plants from three different hybrid genotypes (GT1, GT2, and GT3), (initially grown under control conditions [20°C], then acclimated for one week at 12°C) for 19 days after transfer to cold (4°C) conditions (**Figure 1**). Shoot dry weight (DW), but not fresh weight (FW) continued to increase during the exposure of the plants to 4°C. Consequently, shoot water content gradually decreased by almost half at the end of the recorded time (**Figure 1A**). Simultaneously, FW but also DW of taproots decreased together with the taproot water content during the cold exposure period (**Figure 1A, B**). These results showed that growth of taproots was more affected than that of shoots in the cold and suggested differential physiological and metabolic responses of the shoot and root tissues to cold exposure.

**Figure 1.**
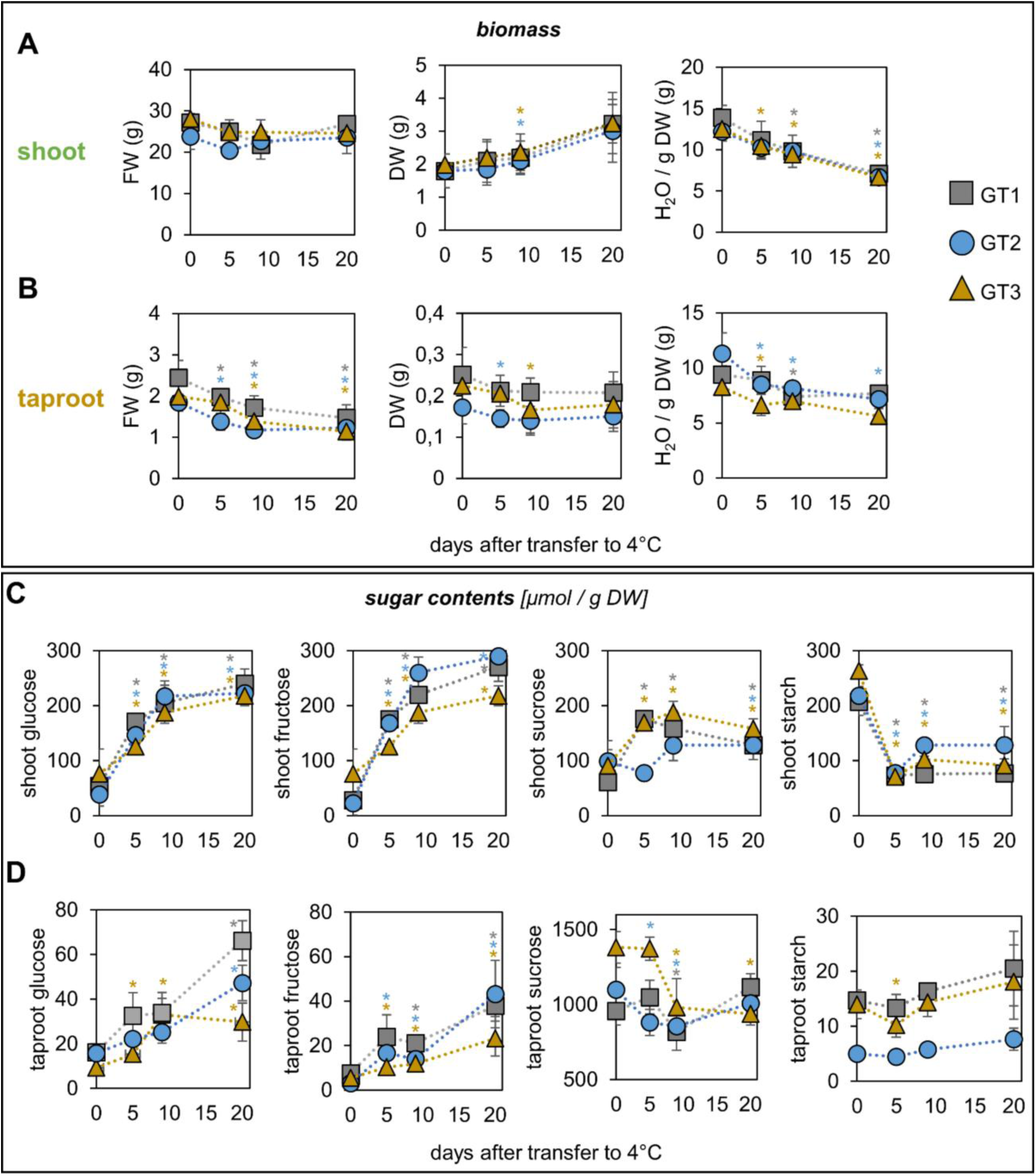
Biomass and sugar accumulation response to cold temperatures in shoots and taproots of 6-week old sugar beet plants from three different genotypes (GT1 = grey square; GT2 = blue circle; GT3 = brown triangle). Plants were grown for six weeks at 20°C, then transferred to 12°C for one week and then to 4°C (start of recording of biomass and sugar accumulation) for 19 days. For each data point, whole organs (shoots or taproots) were harvested at midday. Data points show means from n=6 to 10 plants ± SD. **(A, B)** Fresh weight (FW), dry weight (DW) and water content of shoots and roots. **(C, D)** Sugar and starch accumulation during the course of the chilling (4°C) period in shoots and taproots, respectively. Significant changes to the control condition (first data point) were calculated using double sided Student’s *t*-test (* = *p* < 0.05).

### Sugar levels behave differently in shoots and taproots in the cold

Accumulation of soluble sugars in shoots is a common response to low temperatures and part of cold acclimation process of many plant species (Steponkus, 1971; Wolfe and Bryant, 1999; Strand et al., 1997). Also, in our cold-dependent growth analysis, leaf material (obtained from the very same sugar beet plants as was used for biomass and water content calculation (**Figure 1A**)) exhibited a clear increase in the levels of glucose and fructose (and to a lesser extent of the disaccharide sucrose) after transfer to 4°C (**Figure 1C**). In contrast to soluble sugars, leaf starch contents in all three genotypes decreased rapidly after transfer to 4°C, reaching 20 to 33% of the value present prior to transfer (**Figure 1C, rightmost panel**).

In taproot tissue, sugar accumulation dynamics differed markedly from those in shoots. Glucose and fructose levels slightly increased in the cold, but reached only between 10 to 20 percent of the monosaccharide concentrations of leaves. Prior to transfer to 4°C, taproot sucrose levels exceeded those of monosaccharides 30-to 100-fold. Taproot starch levels of all genotypes were extremely low and did hardly change during cold treatment (**Figure 1D**). The three genotypes analyzed, however, exhibited different sugar and starch accumulation dynamics in the cold. While GT2 and GT3 taproot sucrose levels clearly decreased in the cold, GT1 sucrose levels fluctuated only marginally. Interestingly, the steep drop in sucrose concentration in taproots of GT3 (by about 400 µmol/g DW) and to a lesser extend of GT2 (by about 200 µmol/g DW) was not accompanied by a proportionate increase of monosaccharides, as would be expected for an exclusive hydrolysis of sucrose. These massive losses of taproot sucrose rather suggested that this sugar was either (i) increasingly respired, (ii) converted into compounds other than the monosaccharides glucose and fructose, or (iii) exported from the taproot tissue into other organs. In the following, we aimed to elucidate the fate of sucrose with respect to these possibilities.

### Cold exposure affects photosynthesis rate and carbon dioxide assimilation

In cold tolerant plants like Arabidopsis, sugars accumulate in leaves in the cold when photosynthetic activity is maintained during reduced sucrose phloem loading and increased sugar import into vacuoles of leaf mesophyll cells (Strand et al., 1997; Wingenter et al., 2010; Pommerrenig et al., 2018). We analyzed the impact of cold on sugar beet photosynthesis with pulse amplitude modulated (PAM) fluorometry and CO_2_ assimilation with gas exchange measurements (**Figure 2**). These measurements revealed that Photosystem II quantum yield (Y(II)), leaf CO_2_ concentrations (*C_i_*), CO_2_ assimilation rate (*A*), and leaf transpiration rate (*E*) were dependent on the ambient temperature and that plants exposed to cold responded with a decline in photosynthetic efficiency (**Figure 2**). All three genotypes showed a slight but significant reduction of Y(II) already after one week transfer to 12°C. Simultaneously, non-photochemical quenching Y(NPQ) increased, and non-regulated energy dissipation Y(NO) decreased at this temperature in the leaves of all three genotypes (**Figure 2A**). The higher Y(NPQ) quantum yield at 12 °C compared to 20°C indicated an increased flow of electrons towards the Mehler-Ascorbate peroxidase pathway (Asada et al., 1998) upon exposure to this temperature to undergo e.g. thermal energy dissipation at Photosystem II reaction centers. After transfer to 4°C, Y(II) decreased further and did not recover over the time period tested. However, the decrease of Y(NPQ) quantum yield and the significant increase in Y(NO) quantum yield indicated that electrons underwent unregulated energy dissipation which might induce free radicals and membrane damage at this low temperature (**Figure 2A**). Measurements of CO_2_ gas exchange showed that the reduced PSII activity, as determined by PAM fluorometry was accompanied by a drastic decline of the CO_2_ assimilation rate (*A*) at 4°C but not at 12°C (**Figure 2B**). Transpiration rates (*E*) increased transiently in all three genotypes already at 12°C but more severely at 4°C. The elevated transpiration coincided with a chilling-dependent increase in the leaf CO_2_ concentration, indicating that despite increased stomata opening, activities of Calvin cycle enzymes were greatly reduced (**Figure 2B**).

**Figure 2.**
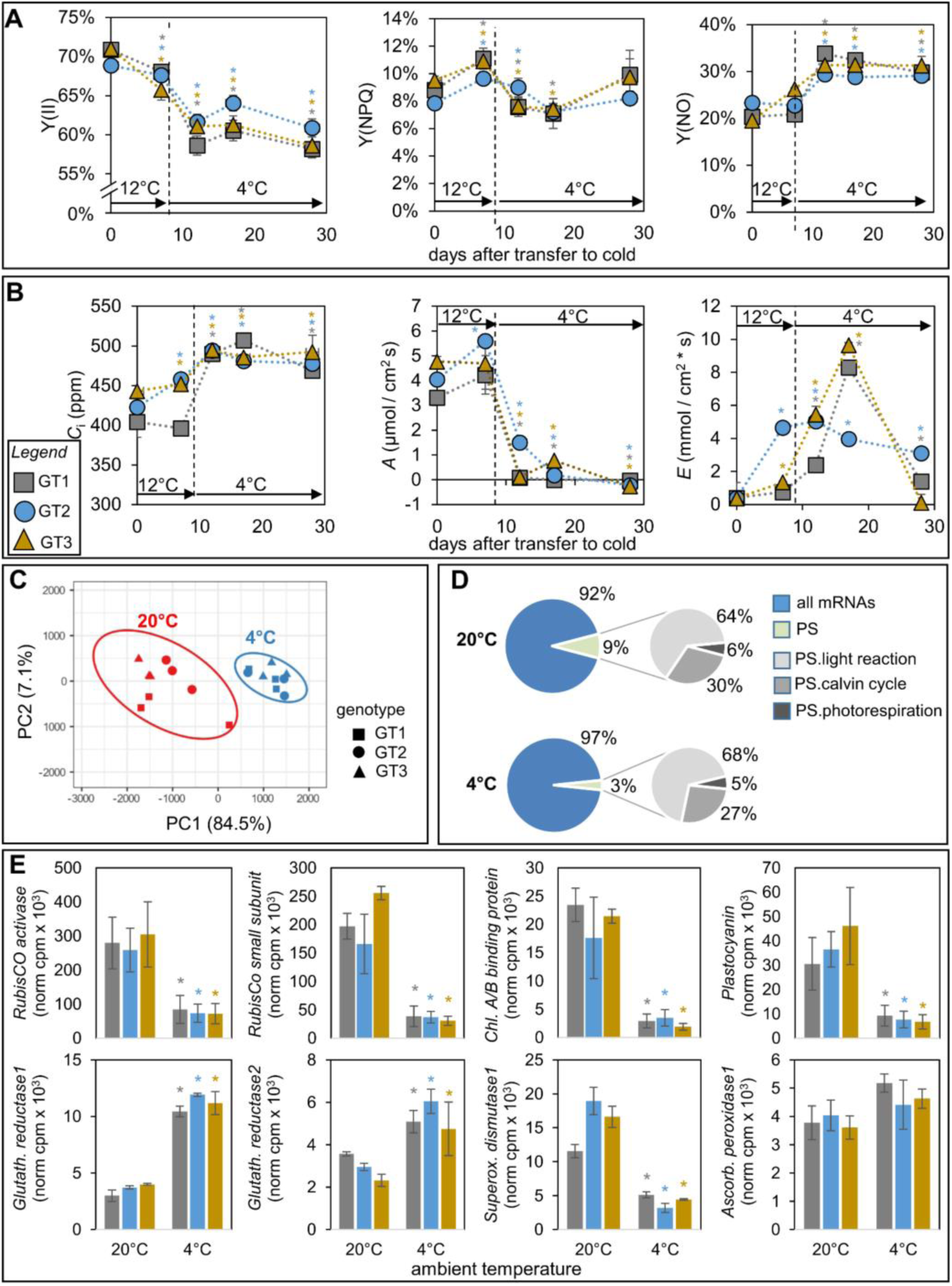
Photosynthetic parameters, CO_2_ assimilation, and expression data of sugar beet leaves after cold exposure. Sugar beet plants of three genotypes (GT1 = grey square; GT2 = blue circle; GT3 = brown triangle) were grown for six weeks at 20°C and then transferred to 12°C for one week and then to 4°C for three weeks. **(A)** PAM measurements of leaves of the three different genotypes. Quantum yield of photosynthesis [Y(II)], of non-photochemical quenching [Y(NPQ)], and of non-regulated quenching [Y(NO)]. At each time point four plants per genotype were analyzed. **(B)** Gas exchange measured for the same plants as used in **A)**. Intercellular leaf CO_2_ concentration (*C*_i_), CO_2_ assimilation rate (*A*), and transpiration rate (*E*) are depicted. For each measurement, four independent plants were used. The very same plants were used for the measurements at the different time points after transfer to cold conditions. Significant changes to the control condition (first data point) were calculated using Student’s *t*-test (* = p < 0.05). **(C)** Principal component analysis (PC1 *versus* PC2) for three genotypes based on expression values of 162 photosynthesis-related genes extracted from RNA-seq data of source leaves from plants grown at 20°C after exposure to 4°C or to control conditions (20°C) for 14 days, respectively. **(D)** Percentage of RNA-Seq reads annotated as genes coding for photosynthesis (PS) related proteins. Pie charts represent the averaged means from three different genotypes at 20°C (control) and after 14 days at 4°C. **(D)** Expression of *RubisCO Activase* (*Bv2_025300_tzou.t1*), *RubisCO small subunit* (*Bv2026840_jycs_t1*), *Chlorophyll A/B binding protein A* (*Bv_002570_dmif.t1*, *Plastocyanin* (*Bv_004160_hgjn.t1*), *Glutathione reductase1* (*Bv3_069540_erom.t1*), *Glutathione reductase2* (*Bv5_120360_jpwm.t1*), *Superoxide dismutase1* (*Bv5_102420_sxsu.t1*), *Ascorbate peroxidase1* (*Bv1_007470_ymzt.t1*). Data represent the mean normalized cpm values of three independent RNA-seq analyses per genotype and temperature condition ± SD. Asterisks represent *p*-values < 0.05 according to double sided *t*-test in comparison to the values at control condition (20°C).

To gain insight into global cold-dependent gene expression of sugar beet source and sink tissues, we performed RNA-seq analyses on leaf and taproot tissue of sugar beet plants from the above genotypes exposed to cold (4°C) or control (20°C) conditions. Samples were collected 14 days after transfer from 12°C to 4°C, i.e. when metabolic accumulation of sugars (**Figure 1**) and photosynthetic rate were maximally contrasting. The obtained RNA-seq reads were mapped to the sugar beet reference genome (Dohm et al., 2013). Transcriptome sequencing data has been deposited in the GenBank Sequence Read Archive (BioProject PRJNA602804).

Exposure to cold induced global rearrangement of gene expression in both shoot and taproot tissues (Supplemental Figure 1). We extracted transcript information on genes involved in photosynthesis. In a PC analysis based on expression values in leaf tissue of genes annotated as ‘photosynthesis’, ‘photosynthesis.lightreaction’, ‘photosynthesis.calvin cycle’, or ‘photosynthesis.photorespiration’ by Mapman Ontology for sugar beet, the PC1 separated the temperature treatments in the three genotypes. PC1 explained 84.5%, PC2 7.1% of the variance in expression between 4°C and 20°C within the genotypes (**Figure 2C**). Independent genotypes were not clearly separated and accordingly, expression levels of photosynthesis-related genes behaved similarly in all three genotypes (**Figure 2C**, Supplemental Figure 1). At 20°C, about 9% of all transcript reads of each genotype could be assigned to ‘photosynthesis’ subgroups. After exposure to 4°C, this group was represented by only 3% of all reads, indicating a drastic downregulation of photosynthesis-related genes in the cold (**Figure 2D**). Downregulation of expression was for example observed for transcripts with homology to genes encoding RubisCO activase (BvRCA), RubisCo small subunit (BvRBCS), a Chlorophyll A/B binding protein (BvCABA), and Plastocyanin (BvPC) (**Figure 2E, upper row**). Genes related to ROS processing on the other hand displayed differential regulation. Whereas genes encoding Glutathione reductases were upregulated in the cold, genes encoding Superoxide-dismutase or Ascorbate reductase were down- or not significantly regulated, respectively (**Figure 2E, bottom row**). In summary, the data demonstrated that sugar beet photosynthesis was extremely sensitive to chilling temperatures below 12°C and suggested that the (hardly occurring) assimilation of CO_2_ does not completely account for the increase in biomass and sugar determined for leaves of cold-treated sugar beet (**Figure 1**).

### Cold temperatures alter major carbohydrate metabolism in shoots and taproots

We investigated whether the reduction of taproot sucrose concentration in the cold could be explained with increased respiration and whether cold conditions would result in differential expression of genes involved in major carbohydrate metabolism **(Figure 3)**. Respiration in taproot tissue was dependent on the examined part of the taproot, in that it decreased with increasing depths of the surrounding soil **(Figure 3A)**. This position-dependent decrease in respiration (proportionate to the depth of soil surrounding the respective part of the taproot) was also observed at 4°C, however, in each part of the taproot, respiration was – in comparison to the corresponding control – generally lower when sugar beets had been exposed to 4°C **(Figure 3A)**. This data suggested that, in the cold, carbohydrates in the taproot were used for glycolytic and oxidative catabolism to a lesser extent than under the 20°C control condition. In shoots, i.e. in source leaves of all genotypes, on the contrary, respiration increased in the cold **(Figure 3B)**, indicating that the mature leaves, which hardly assimilate CO_2_ at this temperature **(Figure 2)**, had a high requirement for carbohydrate supply from other sources. One of these sources was probably starch, which decreased in leaves in the cold (**Figure 1**). PC and heat map analysis, loaded with expression values of genes assigned as “major CHO metabolism”, revealed organ- and temperature-dependent differences **(Figure 3C**, **Figure 3D)**. The first principal component PC1 explained 66.9% of the expression differences between roots and shoots and the PC2 accounted for 17.9% of the differences in expression between 20°C and 4°C. Both organs showed clearer separation at 20°C in comparison to 4°C **(Figure 3C)**. The heat map representation visualizes that expression levels of genes contributing to starch degradation and synthesis in leaves were up- (starch degradation) or downregulated (starch synthesis) by cold exposure, respectively. Despite extremely low starch levels in taproots **(Figure 1)**, starch-related genes were also expressed and regulated in taproots **(Figure 3D)**. This observation is in line with a report from Turesson et al (2014) who showed that starch metabolic enzymes were active despite the lacking occurrence of starch in taproots (Turesson et al., 2014).

**Figure 3.**
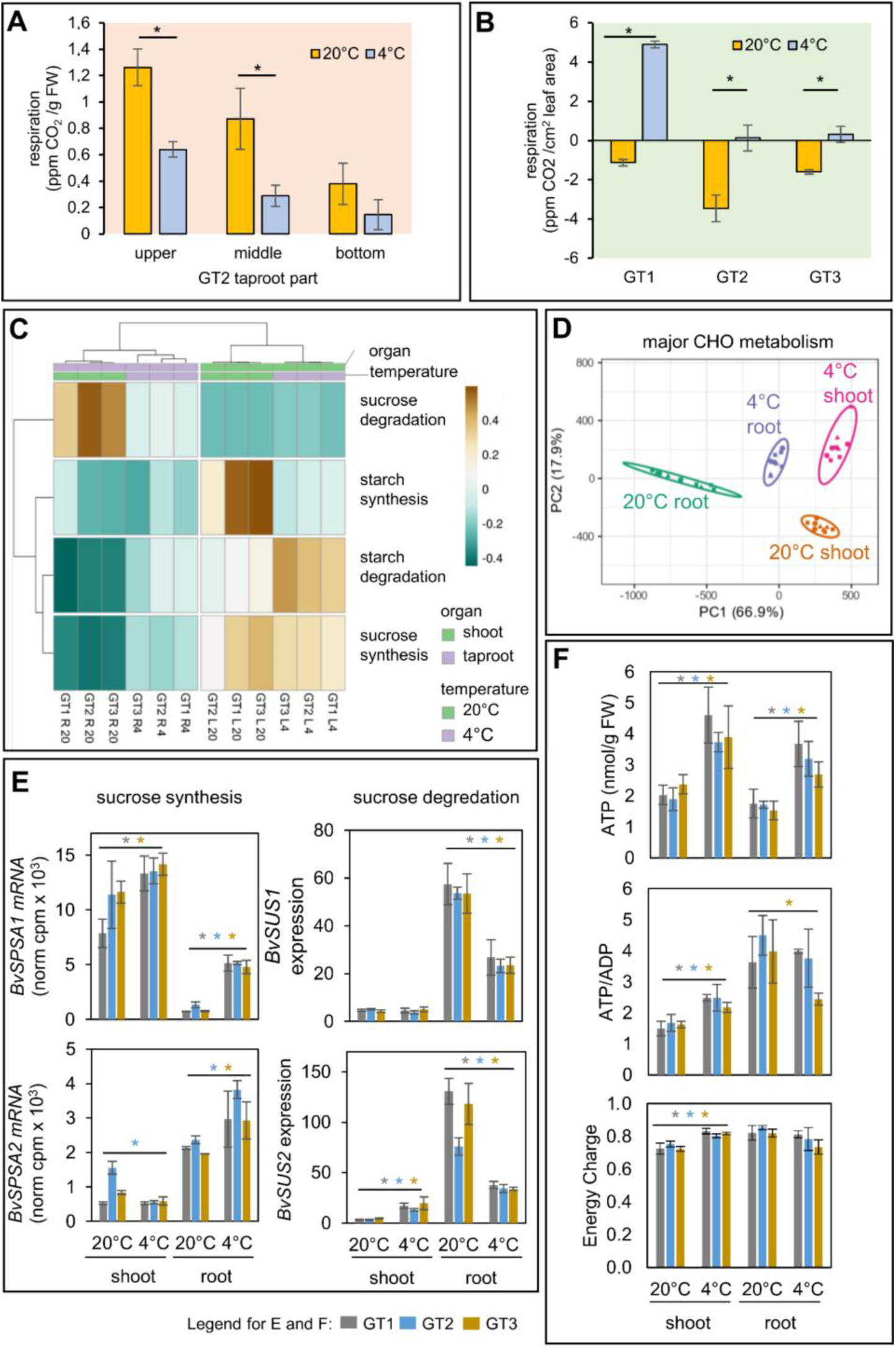
Changes in major carbohydrate metabolism and energy state in response to cold. **(A)** Respiration (CO_2_ production) of different taproot regions from GT1 under control conditions (20°C, yellow bars) or after one week transfer to 4°C (blue bars). **(B)** Respiration (CO_2_ production) from leaf tissue of three genotypes (GT1, GT2, GT3) under control conditions (20°C, yellow bars) or after 1-week transfer to 4°C (blue bars). **(C)** Principal component (PC) analysis (PC1 *versus* PC2) for three genotypes based on expression values of 112 genes with GO annotation “major CHO metabolism” (loadings) extracted from RNA-seq data of source leaves from plants grown at 20°C and transferred for 1 week at 12°C followed by 14 days at 4°C or control conditions (20°C). **(D)** Heatmap analysis of grouped expression values extracted from RNA-seq data. Unit variance scaling was applied to rows. Rows are clustered using Manhattan distance and average linkage. **(E)** Expression values for two Sucrose Phosphate Synthase genes (*BvSPSA1* and *BvSPSA2*) extracted from RNA-seq data of shoots and roots and expression values for two Sucrose Synthase genes (*BvSUS1* and *BvSUS2*) extracted from RNA-seq data of shoots and roots from GT1, GT2, GT3. Data represent the mean normalized cpm values of three independent RNA-seq analyses per genotype and temperature condition ± SD. **(F)** ATP, ATP/ADP ratio, energy charge, EC = [ATP] + 0.5 [ADP]/[ATP] + [ADP] + [AMP]. **(E/F)** Data are means ± SD. Asterisks represent *p*-values < 0.05 according to double sided *t*-test in comparison to the values at control condition (20°C).

Expression levels of sucrose synthesis genes were upregulated in roots in the cold but unchanged in shoots. Sucrose degradation genes, however, were clearly downregulated in roots but slightly upregulated in shoots **(Figure 3D)**. Sucrose Phosphate Synthase (SPS) and Sucrose Synthase (SUS) are key factors of sucrose biosynthesis and degradation and regulate carbohydrate partitioning between source and sink tissues (Voll et al., 2014; Sturm, 1996; Martin et al., 1993; Kovtun and Daie, 1995). A genome-wide search in the sugar beet genome (RefBeet 1.2, (Dohm et al., 2013)) identified two SPS and four SUS isoforms. Bayesian analysis identified both SPS isoforms as homologs of the Arabidopsis SPS ‘A’ subgroup (Voll et al., 2014) (Supplemental Figure 3). The two SPS isoforms showed differential organ-specific and cold-dependent expression. In shoots of all genotypes, expression of *SPSA1* was about 10-fold higher than in roots, when plants had been exposed to 20°C. Cold treatment upregulated its expression in roots up to sevenfold, but did not affect expression levels in the shoot. *SPSA2* expression at 20°C was low in shoots but high in roots of all three tested genotypes. The expression of this isoform was previously identified as taproot-specific, glucose-induced, and sucrose-repressed (Hesse et al., 1995). *SPSA2* expression was also unaltered or even downregulated (in case of GT2) in shoots upon cold treatment, but, as opposed to *SPSA1, SPSA2* expression was induced in taproots of all genotypes. On the protein level, revealed by MS-based analysis of the soluble proteome from the very same taproot tissues as was used for the transcriptome analysis, BvSPSA1 but not BvSPSA2 was slightly upregulated. SPS activity, however, was higher under 4°C in comparison to 20°C in both protein extracts from leaves and taproots (Supplemental Figure 3). Higher levels of UDP in taproots and Sucrose-6-Phosphate in shoots in the cold in comparison to control temperatures along with the elevated levels of the allosteric SPS activator G-6-P (Huber and Huber, 1992) supported a scenario in which SPS activity was elevated in both roots and shoots (Supplemental Figure 3).

The expression of the four sucrose synthase isoforms showed tissue and temperature-dependent differences. While *BvSUS1* and *BvSUS2* isoforms were strongly expressed in roots and their corresponding proteins highly abundant, BvSUS3 and BvSUS4 were hardly expressed and their corresponding proteins were not detected by MS in a soluble proteome fraction (**Figure 3E**, Supplemental Figure 4). Both *BvSUS1* and *BvSUS2* were ten (*BvSUS1*) to hundredfold (*BvSUS2*) higher expressed in roots in comparison to shoots. After the cold exposure period, mRNA levels of both isoforms decreased about half in the roots. *BvSUS2* transcript levels in shoots increased ten to twentyfold, however, without reaching the high levels in taproots (**Figure 3E**). BvSUS2, but not BvSUS1 was also reduced at the protein level indicating differential protein turnover dynamics of the two isoforms in the cold (Supplemental Figure 4).

To determine the cellular energy state of shoot and taproots, adenylate levels were measured **(Figure 3F)**. ATP, ATP/ADP ratio, and energy charge (EC = [ATP] + 0.5 [ADP]/[ATP] + [ADP] + [AMP]) increased in shoots of all genotypes. This elevated energization of shoot tissue in the cold can be explained by the drastic decrease in ATP-consuming CO_2_ assimilation **(Figure 2B)** and the increase of respiration in shoots **(Figure 3B)**. On the other hand, energization of taproot tissue did not change in the cold. Although ATP levels also increased, ATP/ADP ratios of GT1 and GT2 taproots were unaltered or even decreased in GT3. Also EC of taproots did not increase in the cold but rather decreased in tendency in GT2 and GT3 taproots **(Figure 3F)**. Taken together, these data indicate that developing taproots shifted in the cold from a sucrose consuming/storing towards a sucrose synthesizing tissue and that leaves adopted – at least in part – characteristics of sink tissues.

### Cold temperatures reverse phloem translocation of sucrose and esculin

The above data indicated that cold-induced shoot sugar accumulation was not or only insufficiently fueled by carbon dioxide assimilation, or starch degradation, and suggested that carbon used as building block for shoot metabolites might be remobilized from taproot storage cells. To track the fate of taproot-based carbon after exposure to cold temperatures, we directly fed taproot tissue with radiolabeled ^14^C-sucrose by injecting the substance from the exterior into the fleshy parenchymatic taproot tissue of plants grown under 20°C control conditions or cold-exposed plants (5 days at 12°C and then 7 days 4°C). The treated plants were then kept for one more week at control or cold temperatures and then dissected into individual leaves and taproots. The leaves or longitudinal thin sections of taproots were pressed and dried, and incorporated radioactivity was visualized using phosphor imaging plates and software (**Figure 4**, Supplemental Figure 6, Supplemental Figure 7).

**Figure 4.**
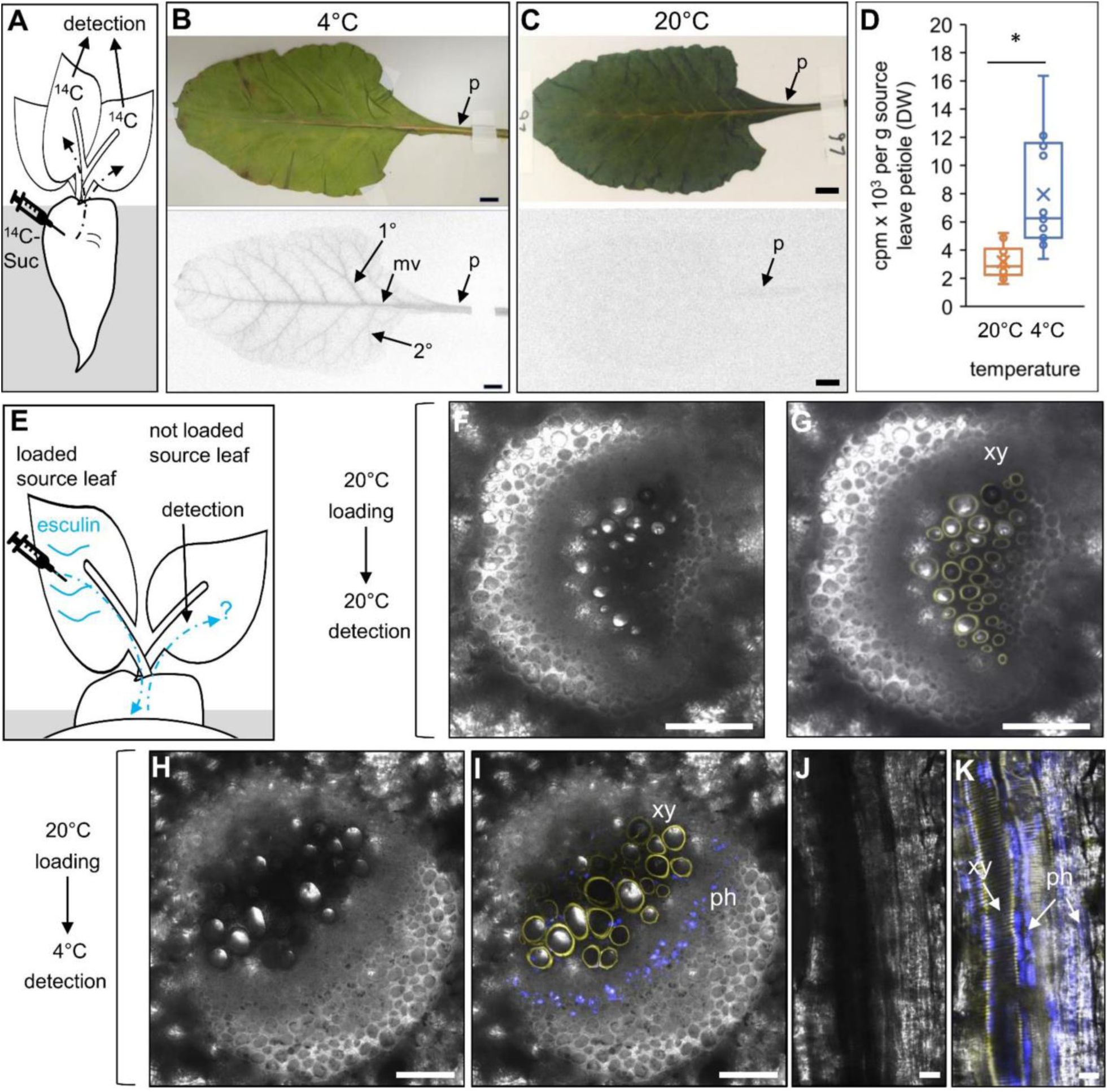
Distribution of ^14^C-sucrose and esculin in leaves. **(A-D)** Autoradiography of ^14^C-sucrose in leaves. **(A)** Schematic depiction of experiment. Taproots were inoculated with ^14^C-sucrose solution and harvested and dried leaves were autoradiographed one week later. **(B)** Source leaf from a representative plant grown for one week under at 4°C. Blackening of veins indicates radioactivity incorporated and distributed into leaf tissue after injection of radiolabeled sucrose into taproots. Abbreviations: p = petiole; mv = middle vein; 1° = first order lateral vein; 2° = second order lateral vein. **(C)** Source leaf from representative control plant grown at 20°C. **(D)** radioactivity in cpm (counts per minute) measured in isolated petioles from plants grown under 4 or 20°C. Center lines show the medians; box limits indicate the 25th and 75th percentiles; whiskers extend 1.5 times the interquartile range from the 25th and 75th percentiles, outliers are represented by dots; crosses represent sample means; n = 16 sample points. **(E-K)** Esculin loadings. Yellow fluorescence indicates lignified xylem vessels, blue fluorescence indicates esculin trafficking. **(E)** Schematic depiction of experiment. Esculin was loaded onto the scratched surface of a source leaf of plants grown at 20°C. Loaded plants were transferred to 4°C or kept at 20°C. Petioles of neighbored, not loaded source leaves were analyzed for esculin fluorescence in plants from 4°C or 20°C. **(F-I)** Cross sections through petiole of a source leaf not loaded with esculin from plants loaded at 20°C. **(F, G)** Petioles from 20°C **(F)** Bright field image. **(G)** UV fluorescence image. **(H, I):** Petioles from 4°C. **(H)** Bright field image. **(I)** UV fluorescence image. **(J, K)** Longitudinal sections of a petiole from 4°C. **J)** Bright field image. **K)** UV fluorescence image. Abbreviations: xy: xylem, ph: phloem. Bars are 50µm in G and H and 100µm in E, F, I, and J.

This analysis surprisingly revealed that plants grown under the 4°C condition showed distribution of radioactivity in source leaves. Radioactivity in leaves of cold-treated plants was detected in leaf veins and intensity gradually decreased towards the leaf tip indicating transport via the phloem vessels (**Figure 4B**). In plants grown under control conditions, however, radioactivity could hardly be detected in source leaves (**Figure 4C**). However, radioactivity was to some extent detectable in young sink leaves of control plants and extractable from combined shoot petioles **(Figure 4D)**. This radioactivity may represent xylem transported sucrose or derivatives due to injury of punctuated vessels as a result of the invasive inoculation procedure. The drastic water loss in shoots upon cold (**Figure 1**) however indicated that at 4°C radiolabeled sucrose was not efficiently transported to prior source leaves via the xylem but rather via the phloem.

To test this hypothesis, we used a strategy less invasive to the organs/tissues examined later, and more realistically mirroring the actual transport of assimilates (including the prior “downward” transport. We loaded esculin, a phloem mobile coumarin glycoside (Knoblauch et al., 2015) recognized by several sucrose transporters, including the *Beta vulgaris* phloem loader BvSUT1 (Nieberl et al., 2017) onto source leaves and assessed esculin transport routes directly via detection of esculin-derived fluorescence in thin sections of leaf petioles of source leaves from the very same plants, which had not been loaded with esculin, after transfer to cold or under control conditions. Here we observed that blue esculin fluorescence was solely detected in phloem of vascular bundles of source leaves from plants transferred to cold. However, the fluorescence was not only confined to the phloem region but also detected to some small extent in a bundle region interspersed with the yellow fluorescence of the lignified xylem vessels **(Figure 4)**. At 20°C, esculin fluorescence was never detected in the phloem (**Figure 4**).

To follow sucrose flow directly from the site of inoculation in the taproots, we performed longitudinal thin sections of taproots inoculated with the radiolabeled sucrose and exposed the tissue to phosphor imaging plates (**Figure 4**, Supplemental Figure 6, Supplemental Figure 7). Radioactivity in taproots from plants exposed to 4°C was detectable and concentrated in veiny or spotty structures that resided between the site of inoculation and the taproot top (crown) tissue. At higher magnification, these structures could be identified as vascular bundles (Supplemental Figure 6). In taproots from plants grown under control conditions, no such distinct darkening of vascular structures could be observed, although some observed blackening of crown tissue indicated that radioactivity was also transported upwards into the direction of the shoot (Supplemental Figure 7). However, in most cases, radioactivity in 20°C taproots was either merely confined to parenchymatic regions near the site of inoculation or concentrated in thick strands that reached from the site of inoculation towards the emergence of lateral roots. These results indicated that radiolabeled sucrose and esculin – the latter first being translocated to the base of the petiole of the loaded leaf and though (at least parts of) the taproot - were preferentially transported from taproots into shoots in the cold but not under control conditions and suggested that sucrose released from parenchymatic storage tissue was also transported in the same manner.

### Vacuolar sucrose importer and exporter genes and proteins show opposite cold-dependent expression

Next, we analyzed whether transport of sucrose from taproots to shoots in the cold could be mediated by differential activity of vacuolar sucrose importers and exporters. In Arabidopsis, vacuolar sucrose import and export are mediated by activity of TST1 and SUC4 transporters, respectively (Schulz et al., 2011; Schneider et al., 2012). In sugar beet, the TST1 homolog BvTST2;1 is responsible for vacuolar sucrose accumulation (Jung et al., 2015). TST2;1 expression in the taproots of all tested genotypes greatly exceeds that in leaf tissue substantiating its role as the sucrose loader of taproot parenchyma vacuoles (**Figure 5**). Interestingly, both mRNA and protein abundance decreased significantly in all genotypes in taproots after cold treatment (**Figure 5B**, Supplemental Figure 8).

**Figure 5.**
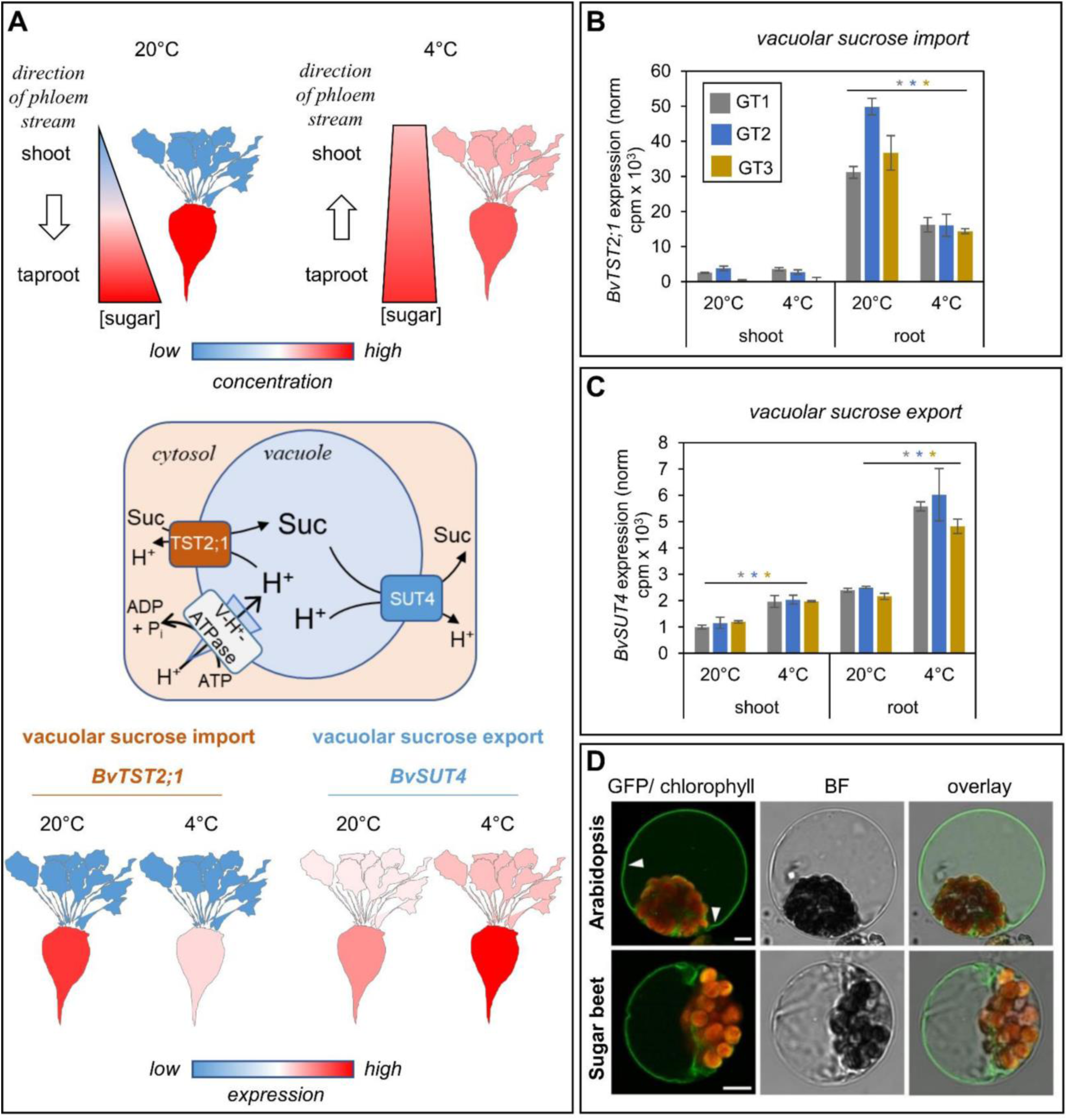
Cold-dependent accumulation of *BvTST2;1* and *BvSUT4* in three different sugar beet genotypes. **(A)** Illustration on cold-induced processes. Upper image: Cold-dependent sugar relocations from taproots to shoots. Middle image: schematic of taproot vacuolar transport processes and factors. Vacuolar ATPase (V-H^+^-ATPase) establishes a proton motif force (pmf) across the vacuolar membrane; TST2;1 acts as proton/sucrose antiporter using pmf for sucrose import into vacuoles. SUT4 acts as proton/sucrose symporter using pmf for vacuolar sucrose export. Bottom image: reciprocal cold-induced regulation of BvTST2;1 and BvSUT4 mRNA levels in taproots **(B)** Transcript abundance of BvTST2;1 (Bv5_115690_zuju) mRNA based on RNA-seq reads. Values represent means from n=3 biological replicates per genotype ± SE. **(C)** Transcript abundance of BvSUT4 (Bv5_124860_zpft.t1) mRNA based on RNA-seq reads. Values represent means from n=3 (mRNA) biological replicates ± SE. Asterisks indicate significant differences between the 20°C and 4°C treatments according to t-test (* = p < 0.05). **(D)** Subcellular localisation of BvSUT4-GFP in Arabidopsis or *Beta vulgaris* leaf mesophyll protoplasts. Single optical sections in all pictures. The green colour shows the GFP-signal; the chlorophyll auto fluorescence is shown in red. Bars = 5 µm. Arrowheads point towards the vacuolar membrane (tonoplast).

Export of sucrose from the vacuole is presumably mediated by a SUC4/SUT4 family homolog. We identified Bv5_124860_zpft.t1 as the unambiguous homolog to the Arabidopsis SUC4 isoform and accordingly termed the corresponding transporter BvSUT4 (Supplemental Figure 10). N-terminal fusions of the BvSUT4 coding sequence with GFP transiently transformed into *Beta vulgaris* or Arabidopsis mesophyll protoplasts clearly indicated that BvSUT4 was a tonoplast located protein (**Figure 5D)**. BvSUT4 mRNA showed lower abundance in older plants in comparison to younger ones (Supplemental Figure 9). In contrast, TST2;1 mRNA increased with progression of leaf development confirming the suggested oppositional activities of the TST2;1 and SUT4 transport proteins (Supplemental Figure 9). In the RNA-seq data from the cold-treated genotypes examined in this study, SUT4 mRNA levels increased significantly in taproots in the cold (**Figure 5C**). These data indicated that vacuolar taproot sucrose import was decreased and vacuolar taproot sucrose release increased under cold conditions and suggested that the opposite regulation of BvTST2;1 and BvSUT4 in taproots was the underlying driving force for the accumulation and delivery of sugars in shoots.

### Expression of floral regulator genes is adjusted in the cold

The observed re-translocation of sucrose from taproots to shoots might represent a preparative metabolic and genetic rearrangement for initiation of flowering. We therefore extracted information on expression of flowering regulator genes and observed significant downregulation of the floral repressor *BvFT1* and upregulation of the floral activator *BvFT2* in the cold in leaves **(Figure 6)**. These results agree with reports from Pin et al (2010), where cold treatment also induced *FT2* and repressed *FT1* expression (Pin et al., 2012). The genotypes analyzed here have biennial growth behavior thus BTC1 and BBX19 may not influence *FT1* expression. However, these two genes were reciprocally cold regulated. While *BTC1* was downregulated in the cold, *BBX19* was upregulated. In contrast to results from Pin et al. (2012), where vernalized biennials had increased *BTC1* mRNA levels in comparison to non-vernalized plants (Pin et al., 2012), BTC1 was downregulated in the cold. However, in the mentioned study, expression was analyzed after and not during early stages of vernalization. We found that *BTC1* and *BBX19* were expressed in both, shoots and taproots, and expression of *BBX19* in taproots exceeded that in the shoot at 20°C almost threefold. However, potential targets of these encoded loss-of-function proteins, *FT1* and *FT2* were specifically and exclusively expressed in leaf tissue (**Figure 6**). In summary, these data showed that the vernalization process was already transmitted to the expression level of floral regulator genes and that transcriptional changes of related genes did occur in both, shoots and taproots.

**Figure 6.**
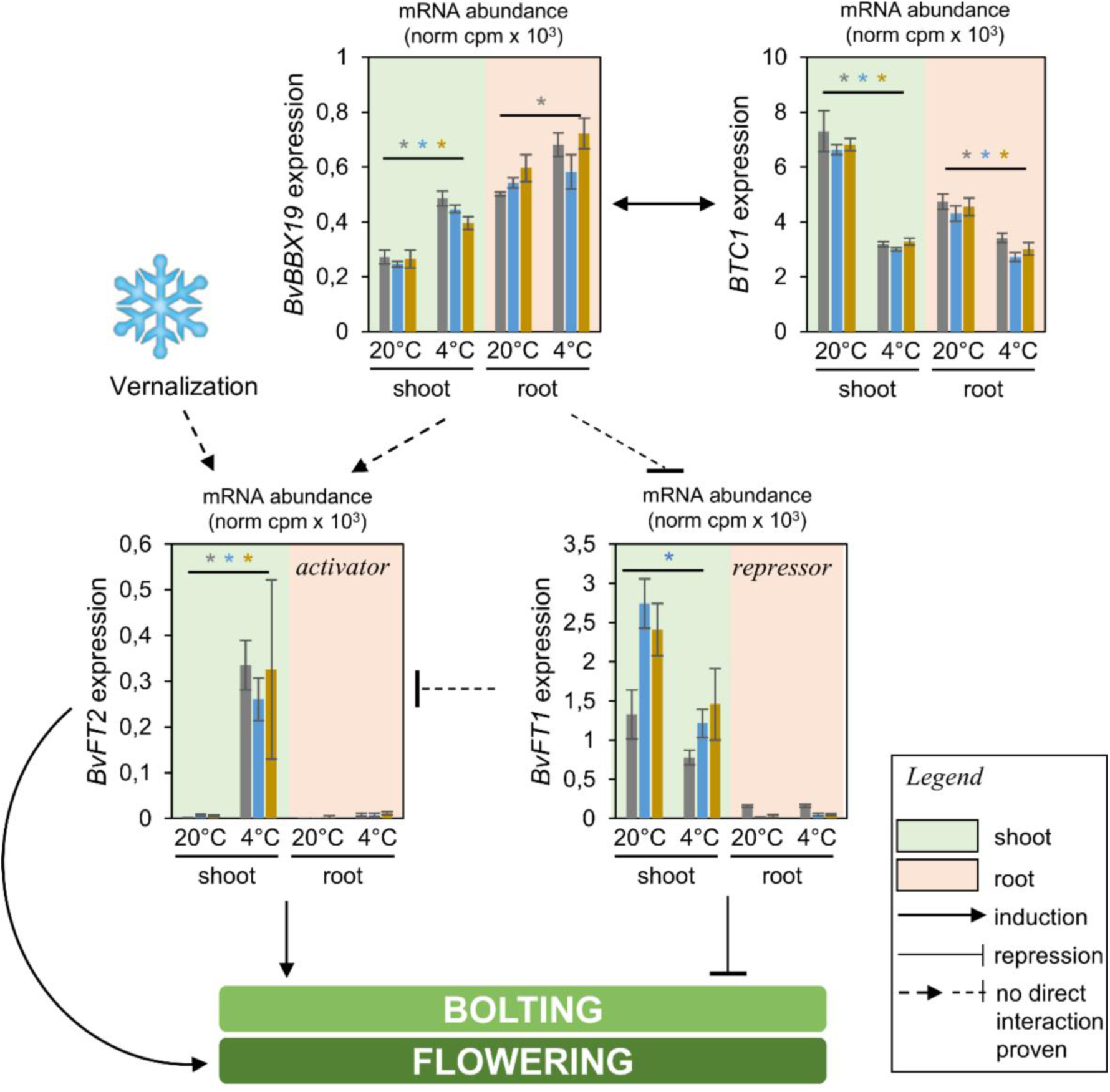
Expression of floral regulator genes. Transcript abundances of *BvBBX19* (Bv9_216430_rwmw.t1), *BvBTC1* (Bv2_045920_gycn.t1), *BvFT1* (Bv9_214250_miuf.t1), and *BvFT2* (Bv4_074700_eewx.t1) based on RNA-seq reads in shoots and taproots of three different genotypes. Values represent means from n=3 biological replicates ± SE. Asterisks indicate *p*-values < 0.05 according to double sided *t*-test.

## Discussion

In this work we discovered a so far unknown switch of sink and source identities of taproots and shoots upon cold exposure of sugar beet plants. In contrast to sinks like seeds, culms or tubers, which adopt source identities after complete differentiation and subsequent separation from their nourishing source, the sink-source switch in sugar beet occurred in response to an environmental stimulus when both shoot and taproot tissues were still physiologically connected.

At 4°C, shoot CO_2_ assimilation was drastically reduced but PSII activity stayed relatively high **(Figure 2).** This correlation indicates that enzymes of the Calvin-Benson cycle slowed down in the cold and could not utilize electrons liberated from the photosynthetic electron transport (PET) chain. During the decreased CO_2_ fixation rates at cold conditions, high PET rates may have detrimental effects because they produce harmful reactive oxygen species (ROS) like super-oxide, hydrogen peroxide or hydroxyl anions (Suzuki and Mittler, 2006; Choudhury et al., 2017; Pommerrenig et al., 2018). During the cold exposure kinetic (**Figure 1 and Figure 2**), we recorded decreased CO_2_ assimilation already at 12°C. As indicated by the increased Y(NPQ) percentage (**Figure 2A**), the Mehler-Ascorbate Pathway (Asada, 1999) might act as an additional quencher for PET-released electrons at this temperature. At 4°C, this scavenging pathway apparently also slowed down, as indicated by the further decrease in Y(II) but also Y(NPQ), and the concomitant increase in Y(NO). The significant increase in Y(NO) at 4°C is indicative for non-regulated energy dissipation, which can severely damage chloroplast membranes and plant cells in a cold- and high light dependent manner. Under those sustained challenging conditions, the cold response was apparently transduced to the level of gene expression where it led to an effective downregulation of transcripts of photosynthesis-related genes **(Figure 2D and 2E)**. Induction of glutathione reductase genes supported a scenario in which leaves induced cellular counter measures against light-induced electron overflow at the photosystems and damages of chloroplasts **(Figure 2E)**.

These data are in agreement with results from Arabidopsis, where photosynthesis as well as expression of *RBCS* and *CAB* genes were significantly reduced after shifting of 23°C-grown plants to 5°C, although photosynthesis recovered after prolonged exposure to cold (Strand et al., 1997). In summary, these metabolic and transcriptomic changes would eventually result in drastic decrease of CO_2_ incorporation into sugars, which are required for growth and protection of cell vitality in the cold.

Despite inactivation of photosynthesis, however, sugars continued to accumulate in leaves and decreased in taproots in the cold **(Figure 1)**. Decreasing sucrose levels in taproots and impaired respiration in root tissue indicated that sucrose was not used for energy metabolism during cold at the same rate as under control conditions in the taproot **(Figure 3)**. Cold tolerant plants like Arabidopsis accumulate sugars in leaves in the cold by maintaining photosynthetic activity, reducing sucrose phloem loading, and increasing sugar import into leaf vacuoles (Wingenter et al., 2010; Nägele and Heyer, 2013). While Arabidopsis has the ability to overcome sugar repression of photosynthesis after prolonged exposure to cold (Huner et al., 1993; Strand et al., 1997), such mechanism apparently does not occur in sugar beet in the same manner. In contrast, the drastic decrease of photosynthetic activity in the shoot rather turned leaves into sink organs, which were supplied with sugar from taproots **(Figure 4)**.

Under non-chilling temperatures, reversibility of taproot sink and remobilization of sugars from storage vacuoles might become essential when leaves have to re-grow after wounding of the shoot caused by e.g. feeding damage or when a new strong sink like the inflorescence is formed after winter. However, as indicated by the movement of radiolabeled sucrose and fluorescent esculin towards the shoot, and therefore to previous source leaves also early cold response triggered a remobilization of carbohydrates from taproot storage **(Figure 4)**. While sucrose biosynthesis and hydrolysis were reciprocally regulated under warm and cold conditions **(Figure 3)**, levels of taproot sucrose decreased upon cold treatment (**Figure 1**). In agreement with this process, we identified opposing regulation of the major vacuolar sucrose importer (BvTST2;1) and putative major exporter (BvSUT4) in the same tissue (**Figure 5**). BvTST2;1 expression and protein abundance was significantly downregulated, while, in contrast, BvSUT4 was upregulated in the cold. The role of BvSUT4 as an exporter of sucrose is supported by its general homology to sucrose transporters of the SUC/SUT family and by its homology to AtSUC4 (**Figure 5**), for which both sucrose export activity and vacuolar localization have been shown (Schulz et al., 2011; Schneider et al., 2012). It seems unlikely that sugars are released from vacuoles in the cold as monosaccharides via other transporters, e.g. by the already described BvIMP protein (Klemens et al., 2014). This is because vacuolar invertases - a prerequisite for vacuolar monosaccharide generation and thus export – are hardly active at the analyzed developmental stage (Giaquinta, 1979; Godt and Roitsch, 2006).

The previously explained findings are schematically explained in the following model **(Figure 7)**. It is surprising that flux transition occurred already pre-bolting i.e. before the formation of an inflorescence that would then act as new sink organ utilizing remobilized taproot sugars as building blocks. During the early phases of vernalization warmer temperatures or longer daylight, additional prerequisites for bolting (Mutasa-Göttgens et al., 2010; Ritz et al., 2010), do not yet signal onset of spring. However, simultaneously to the switching of identities, the cold exposure also led to adjustment of expression levels of floral regulator genes. *FT1* and *FT2*, the floral repressor and activator genes (Pin et al., 2010), respectively, showed reciprocal regulation in the cold in shoots **(Figure 6)**. The expression of the flowering-related genes *BTC1* and *BBX19* in taproot tissue suggested that taproots might also be involved in the perception of vernalization. It is tempting to think into a direction where newly identified (Pfeiffer et al., 2014; Broccanello et al., 2015; Tränkner et al., 2017) or yet undiscovered bolting loci might harbor yet uncharacterized factors which might integrate both, bolting and required sink-source transition, similar to the recently described FT homolog StSPS6A (‘tuberigen’) in potato (Navarro et al., 2011; Abelenda et al., 2019).

**Figure 7.**
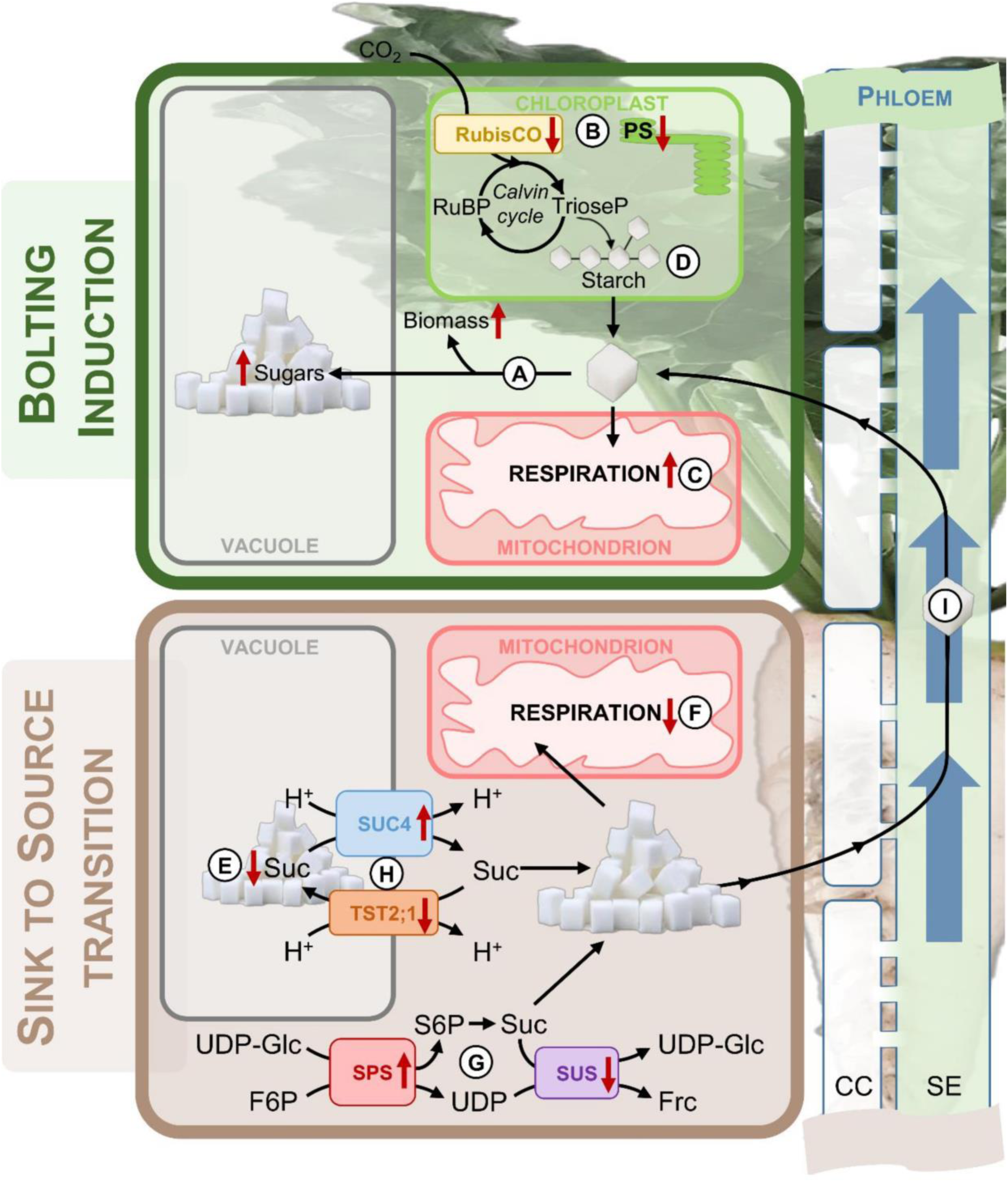
Schematic illustration of cold-induced sink to source transition. Leaf- and taproot-tissue of sugar beet are reprogrammed and source and sink identities shifted upon cold. Shoots adopt sink identity during cold. Biomass and sugar concentration in the shoot increase **(A)** despite reduced photosynthetic activity and inactivation of carbon assimilation **(B)**. Concomitantly, shoot respiration increases **(C)** and cellular starch pools decrease **(D)**. Contrastingly, taproots show a decrease of sucrose levels **(E)** but lower respiration rate **(F)** as well as increased sucrose biosynthesis **(G)**. Taproot sugar is remobilized in the cold due to opposite regulation of taproot-specific vacuolar sucrose importer (BvTST2;1) and exporter (BvSUT4) activity **(H)**. Taken together, this results in a reversal of the phloem translocation stream **(I)** triggered by a reprogramming of source and sink identities, which might correlate with inflorescence initiation.

Our study represents a comprehensive analysis of sugar beet taproot tissue during cold treatment and shows that cold temperatures induce a sink to source transition, which establishes accumulation of taproot-based carbohydrates in the shoot. For this, sugars have to be loaded into taproot phloem, transported from taproots to shoots, and unloaded in leaf tissue. Currently it is unknown whether taproot phloem loading in the cold involves an apoplastic step, whether the same phloem vessels are being used for root- and shoot-bound sugar trafficking, and how sugar unloading is established in former source leaves. Latter issue possibly involves a reprogramming of transporter activity that could mediate sugar efflux from the vasculature to the mesophyll involving both passive and active transport processes. Our transcriptomic and proteomic approach might reveal candidate factors and transporters involved in this unloading in the cold in the future.

The findings also have implications for agriculture and breeding, where attempts have been made to grow sugar beet over all seasons (Hoffmann and Kluge-Severin, 2011; Hoffmann and Kenter, 2018), a scenario which will become more and more realistic by the generation and employment of bolting-resistant hybrid genotypes (Pin et al., 2010; Pfeiffer et al., 2014; Tränkner et al., 2016). In addition, biennial growth of sugar beet might become facilitated by the increasing occurrence of climate change-induced “warm” winters in e.g. middle and Northern Europe (Lavalle et al., 2009) that would allow cultivation of sugar beet under non-freezing, non-lethal low temperatures.

However, even under non-freezing, but prolonged above zero chilling conditions, the advantages of a longer vegetation period would be negated, at least to some extent, by the herein described trade-off of cold-induced taproot sugar loss. This phenomenon might also partially account for the observed reduced yield and higher marc to sugar ratio of autumn- or early spring-sown sugar beet plants (Hoffmann and Kluge-Severin, 2011).

In future, it will be highly valuable to analyze this observed sink-source transition of taproots in bolting resistant mutants without the activating function of FT2 (Pin et al., 2010) to reveal whether FT activity is required for triggering this transition. Equally relevant will be the generation of BvSUT4 mutant plants to study effects of lacking vacuolar sucrose efflux for floral induction and cold tolerance. Such modified plants would possibly exhibit a diminished taproot sucrose release and therefore a reduced building block supply for inflorescence formation. This potential impact on bolting makes BvSUT4 a highly relevant target for breeding approaches (Pfeiffer et al., 2014; Chiurugwi et al., 2013) aiming at bolting resistance and at withholding cold-induced sucrose loss from taproots.

## Materials and Methods

### Plant Material and Growth conditions

Three hybrid sugar beet genotypes (GT1, GT2, GT3; KWS SAAT SE, Germany) were used for this study. Plants were germinated and grown on standard soil substrate ED73 (Einheitserdwerke Patzer, Germany)/ 10% (v/v) sand mixture under a 10 h light/14 h dark regimen, 60% relative humidity, and 110 µmol m^-2^ s^-1^ light intensity. For growth- and sugar accumulation kinetics, plants were grown for 6 weeks at 20°C, transferred for 1 week at 12°C and then 3 weeks at 4°C. For RNA-seq and proteome analysis, plants were grown for 10 weeks at 20°C, transferred for 1 week at 12°C and then 2 weeks at 4°C. Control plants were kept at 20°C. For harvest, plants were dissected into shoot and taproot tissues. 4 pools out of three different plants were made for each tissue. Tissues were chopped with a kitchen knife, transferred to liquid nitrogen, and kept at −80°C until further processing.

### Chlorophyll Fluorescence Measurements

Photosynthetic activity was measured using an Imaging-PAM *M-Series*-System (Heinz Walz, Effeltrich, Germany). Plants were placed in the dark for 12 min to deplete the energy of PSII. Capacity of PSII was measured by saturation with 14 cycles of PAR 76 (µmol photons m^-2^ s^-1^) light-pulses at 0s, 50s, and 70s. Recorded fluorescence was used for calculation of effective quantum yield of PSII [*Y(II) = (Fm’-F)/Fm’*], quantum yield of non-photochemical quenching [*Y(NPQ) = 1 - Y(II) - 1/(NPQ+1+qL(Fm/Fo-1))*] and of non-regulated energy dissipation [*Y(NO) = 1/(NPQ+1+qL(Fm/Fo-1))*]. Required factors were calculated by the formulas [*NPQ = (Fm-Fm’)/Fm’*], [*qN = (Fm-Fm’)/(Fm-Fo’)*], [*Fo’ = Fo/ (Fv/Fm + Fo/Fm’)*], [*qP = (Fm’-F)/(Fm’-Fo’)*] and [*qL = (Fm’-F)/(Fm’-Fo’) x Fo’/F = qP x Fo’/F*].

### Gas Exchange Measurements

A GFS-3000 system (Heinz Walz, Effeltrich, Germany) was employed to analyze gas exchange-related parameters. A 2.5 cm^2^ gas exchange cuvette was used to measure CO_2_-assimilation rate, respiration, leaf CO_2_ concentration, and transpiration of sugar beet source leaf. Leaf regions including large central mid ribs were omitted. The conditions inside of the cuvette were set to the same temperature, humidity and CO_2_-concentration the plants had been grown at. Measurement sequence is listed in **Table 1**. The listed intervals were determined by a trial-experiment, in which the time necessary for stabilization of the flow of CO_2_ after transfer of the leaf section into the cuvette and adoption to the changed light-intensities was measured. The measurement was started after stabilization of the CO_2_-flow, which required about 5 minutes. Measurements were performed with 4 plants in 3 technical (repeated measurements of the same plant) replicates over a time of 1 min for each condition to account for variation caused by observed natural leaf-fluctuation and leaf area outside of the cuvette. The 30 second interval between the measurements was necessary for the leaf to return to the stabilized value.

**Table 1:** Sequence for gas-exchange measurements.

| time [s] | Light-intensity | measurement |
| --- | --- | --- |
| +0 | PAR 0 |  |
| +220 | PAR 0 | photosynthetic activity |
| +30 | PAR 0 | photosynthetic activity |
| +30 | PAR 0 | photosynthetic activity |
| +460 | PAR 125 | respiration/transpiration (light) |
| +30 | PAR 125 | respiration/transpiration (light) |
| +30 | PAR 125 | respiration/transpiration (light) |
| +320 | PAR 0 | respiration/transpiration (dark) |
| +30 | PAR 0 | respiration/transpiration (dark) |
| +30 | PAR 0 | respiration/transpiration (dark) |

### Respiration of sugar beet taproot tissue

Respiration of taproots was measured by cutting out 0.5 cm^2^ tissue cubes from central taproot regions and measuring CO_2_ production in a whole-plant cuvette with a volume of 60 cm^3^. Values were normalized to tissue weight.

### RNA extraction and sequencing

RNA was isolated from three biological replicates per genotype, tissue (leaf and root, respectively) and treatment, respectively. About 100 mg frozen plant material were pulverized in a tissue lyser (Qiagen, Hilden, Germany) at 30 Hz for 90 sec. After grinding, samples were again transferred to liquid N_2_, supplemented with 1.5 ml QIAzol Lysis reagent (Qiagen, Hilden, Germany), vortexed three times for 30 sec, and centrifuged at 4 °C for 10 min at 12,000 *g*. Supernatants were transferred to fresh tubes, incubated at room temperature (RT) for 5 min, extracted with 300 µl chloroform, vortexed for 15 sec, and centrifuged at 4 °C for 15 min at 12,000 *g*. Aqueous supernatants were transferred to fresh tubes and RNA precipitated with 750 µl isopropanol for 10 min at RT and spun down at 4 °C for 10 min at 12,000 *g.* Precipitates were washed with 75% EtOH and the RNA pellets dried at 37 °C for 5-10 min prior to resuspension in 100 µl DEPC-H_2_O by gentle shaking at 37 °C for 5-10 min. To remove residual contaminants, RNA was further purified using the RNeasy KIT (Qiagen, Hilden, Germany). Per 100 µL RNA suspension, 350 µl RLT buffer (provided with the kit) were added and vortexed briefly. Then, 250 µl ethanol were added and the mixture was vortexed again. The RNA was spin-column purified and finally eluted from the column for a final volume of 50 µl (in DEPC-H_2_O) per sample. The RNA was quantified (NanoDrop 2000/2000c, Thermo Fisher) for each sample prior to further processing or storage at −80 °C. RNA quality was confirmed using an Agilent Technologies 2100 Bioanalyzer (Palo Alto, CA, USA). RNAs (2 µg per sample) were transcribed to cDNAs and sequenced using an Illumina, Inc. HiSeq 2000 system. Sequencing and assembly were provided as a custom service (GATC GmbH, Konstanz, Germany). The statistical analysis process included data normalization, graphical exploration of raw and normalized data, test for differential expression for each feature between the conditions and raw *p*-value adjustment. The analysis was performed using the R software (Team, 2017), Bioconductor (Gentleman et al., 2004) packages including DESeq2 (Anders and Huber, 2010; Love et al., 2014) and the SARTools package developed at PF2 – Institute Pasteur.

### Phylogenetic analysis

Multiple sequence alignments of amino acid sequences were performed using Clustal Omega (Sievers et al., 2011). Bayesian phylogenetic analysis was performed with MrBayes version 3.2 (Ronquist et al., 2012). MrBayes always selected the best-fit models ‘Jones’ (Jones et al., 1992) and ‘WAG’ (Whelan and Goldman, 2001) for amino acid substitution analysis of SPS proteins and SUS proteins, respectively. MrBayes conducted two parallel Metropolis coupled Monte Carlo Markov chain analysis with four chains for 300,000 generations. Trees were sampled every 1,000 generations. The analyses were run until the standard deviation of split frequencies were below 0.005. Consensus trees were computed after burn-in of the first 25% of trees and visualized using FigTree version 1.4.3.

### PCA and heatmap analysis

For RNAseq data the mean cpm values were used for the analysis. Data were visualized using ClustVis (Metsalu and Vilo, 2015).

### Analysis of soluble sugars and starch

Leaves and taproots were harvested separately, frozen in liquid nitrogen, freeze-dried and stored at −80°C until use. Pulverized material was extracted twice with 1 ml 80% EtOH at 80°C for 1 h. Combined extracts were evaporated in a vacufuge concentrator (Eppendorf, Hamburg, Germany) and pellets were resolved in ddH_2_O. For starch isolation pellets were washed with 80% EtOH and 1 ml ddH_2_0. 200 µl water were added to the pellet and the sample was autoclaved for 40 min at 121°C. 200 µl enzyme-mix (5 U α-Amylase; 5 U Amyloglucosidase in 200 mM Sodium-Acetate pH 4.8) were added to the pellet and starch was hydrolytically cleaved into glucose-units at 37°C for 4 h. The enzymatic digestion was stopped by heating the samples to 95°C for 10 min. After centrifugation (20,000 *g*; 10 min; 21°C) the supernatant could be used for starch quantification. Extracted sugars and hydrolytically cleaved starch were quantified using a NAD+-coupled enzymatic assay (Stitt et al., 1989).

### Analysis of phosphorylated metabolites

The contents of phosphorylated intermediates (Glucose-6-Phosphate, Fructose-6-Phosphate, Sucrose-6-Phosphate, UDP-Glucose, UDP) were determined according to (Horst et al., 2010).

### Radiolabeled sucrose translocation assay

Ten- to 12-week old sugar beet plants grown at 20°C under short day conditions (10 h light, 14 h darkness) were used for the analysis. Plants for cold-treatment were grown for 1 more week at 12°C and then kept for 6 to 7 days at 4°C. Taproots from 4°C and 20°C plants were partially uncovered from surrounding soil substrate and a 1 mm hole punched with a biopsy stance into the upper half of the taproot (approximately 1 cm below the soil surface). The created pit was filled with 10 µl of 1 to 2 diluted radiolabeled sucrose (536 mCi/mmol) (Hartmann Analytic, Braunschweig, Germany) and coated with a drop of Vaseline. Plants were then kept for another 10 days at 4°C or 20°C (control). At the end of the treatment, all source leaves of injected plants were detached and individually pressed between blotting paper. For detection of radioactivity in taproots, taproots were dug out, washed and cut in thin slices (approximately 0.5 mm thick) with a truffle slicer and pressed between blotting paper. Radioactivity was recorded with Phosphor-Image plates (exposed for 4 to 5 h to adaxial surface of pressed and dried leaves or to dried taproot slices) and plates were analyzed with a Cyclone Storage Phosphor Screen (Packard Bioscience, Meriden, CT, USA). For quantification of radioactivity in petioles, source leaf petioles from the same leaves used for phosphoimaging were cut off, ground, and pulverized. 5 to 10 mg powder were mixed with 2 ml scintillation cocktail and counts per minute (cpm) recorded with a TRI-Carb 2810TR liquid scintillation analyzer (Perkin Elmer, Waltham, MA, USA).

### *In planta* esculin transport

Ten-week old sugar beet plants grown at 20°C under short day conditions (10 h light, 14 h darkness) were used for the analysis. One source leaf per plant (usually from leaf stage 10 to 12) was abraded at the adaxial side with fine sandpaper (grade 800). About 500 µl of a 100 mM esculin sesquihydrate (Carl Roth, Karlsruhe, Germany) solution was distributed over the injured leaf surface with a plastic pipette. Treated leaves were coated with plastic foil, kept for 2 more days at 20°C and then transferred to 4°C or kept at 20°C (control). After 5 to 7 days in the cold, not esculin-loaded source leaves were detached and sections of petioles were analyzed for esculin fluorescence with a Leica TCS SP5II confocal microscope (Leica, Mannheim, Germany) using a HCX PL APO lamda blue 20.0×0.70 IMM UV objective. Slices of taproots from the very same plants were analyzed for esculin fluorescence to ensure that esculin was successfully translocated into taproots in both cold-treated and control plants. The emission bandwidths were 440 – 465 nm for detection of esculin fluorescence and 594 – 631 nm for lignin fluorescence.

### Soluble protein extraction

Plants were harvested, washed, and separated in the cold into taproots and source leaves. Frozen leaf-tissue was pulverized with N_2_(l) using a Retsch mill (Retsch GmbH, Germany). 800 µl buffer E1 (50 mM HEPES-KOH pH 7.5, 10 mM MgCl_2_, 1 mM EDTA pH 7.5, 2 mM DTT, 1 mM PMSF, 1 mM Pefabloc, 5 mM aminohexanoic acid, 0,1% (v/v) Triton X-100, 10% (v/v) glycerol) were transferred to 100 mg of pulverized tissue into 1.5 ml Eppendorf cups. Samples were vortexed and centrifuged for 3 min at 12.000g at 4°C. 500 µL of the supernatant were loaded onto a Sephadex NAP5 (G25) column (GE Health Care, United Kingdom), pre-equilibrated with buffer E1 w/o Triton X-100. Eluents were collected in precooled Eppendorf cups and stored at −20°C. Taproot tissues were treated as above with the following alterations: Taproots were blended with buffer E1 at 4°C until a homogenous pulp was obtained. The pulp was roughly filtered through a kitchen sieve and centrifuged. 5 ml of the supernatant were dialyzed trough a membrane with 12 kDa pore size for 48 h against 2 L ddH_2_O. Water was exchanged seven to eight times. Samples were collected in precooled Eppendorf cups and used for enzymatic tests or stored at −20°C. Liquid chromatography and tandem mass spectrometry was performed as described in (Jung et al., 2015).

### Isolation of taproot vacuoles and vacuolar proteins

Vacuoles were isolated as described by (Jung et al., 2015).

### Sucrose Phosphate Synthase assay

80 µg of soluble protein were added to 200 µl freshly prepared E_max_ (50 mM HEPES-KOH pH 7.5, 20 mM KCl, 4 mM MgCl_2_, 12 mM UDP-Glc, 10 mM Frc-6-P: Glc-6-P (1:4)), E_lim_ (50 mM HEPES-KOH pH 7.5, 20 mM KCl, 4 mM MgCl_2_, 4mM UDP-Glc, 2mM Frc-6-P: Glc-6-P (1:4), 5 mM KH_2_PO_4_) and E_blank_ (= E_max_ w/o UDP-glucose and sugar-phosphates), respectively. Samples were incubated for 20 min at 25°C, followed by 5 min at 95°C to stop the reaction and centrifuged at 12.000 *g* at 4°C for 5min. 100 µL of the supernatant were pipetted to 100 µL 5 M KOH and incubated 10 min at 95°C. The solution was mixed with 800 µL anthrone (14.6 M H_2_SO_4_, 0,14% (w/v) anthrone) and absorbance immediately measured at 620 nm. A calibration-standard was made with 0-5 mmol sucrose.

### Subcellular localization of BvSUT4 in Arabidopsis and sugar beet mesophyll protoplasts

The BvSUT4 CDS (Bv5_124860_zpft.t1= BVRB_5g124860) was amplified from *B. vulgaris* leaf RNA with the gene specific primers BvSUT4-CACC-f (5’-CAC CAT GAC AGG CCA GGA CCA AAA TA-3’) and BvSUT4-rev (5’-TAC ATG CAT CAC ATG AAC TCT GG-3’). The resulting open reading frame was cloned into pENTR/D-TOPO (Life Technologies, Darmstadt, Germany), sequenced and recombined into the Gateway-compatible destination vector pK7FWG,0 (Karimi et al., 2002) to obtain a p*35S::BvSUT4-GFP* fusion. Transient transformation of A. thaliana mesophyll protoplasts was performed as described (Abel and Theologis, 1994). Isolation and transient transformation of *B. vulgaris* mesophyll protoplasts were performed as described (Nieberl et al., 2017).

### Data availability

Transcriptome sequencing data has been deposited in the GenBank Sequence Read Archive under BioProject PRJNA602804.

## Acknowledgements

The authors would like to thank

Michaela Brock, David Pscheidt (both FAU Erlangen-Nürnberg), Tim Seibel (University of Kaiserslautern) for excellent technical assistance and Karin Fiedler (KWS SAAT SE) for provision of *Beta vulgaris* seed material and management of sugar beet growth.

## Author contributions

H.E.N., F.L., W.K., K.H., U.S., B.P., designed the research;

C.M.R., C.M., I.K., W.Z., F.R., P.N., B.P performed research;

O.C., J.M.C.G, T.M. contributed new analytic/computational/etc. tools;

C.M.R., C.M., B.P., analyzed data;

B.P., C.M., and H.E.N. wrote the paper.

