## Supplementary information for "Vernalization alters sugar beet (*Beta vulgaris*) sink and source identities and reverses phloem translocation from taproots to shoots"

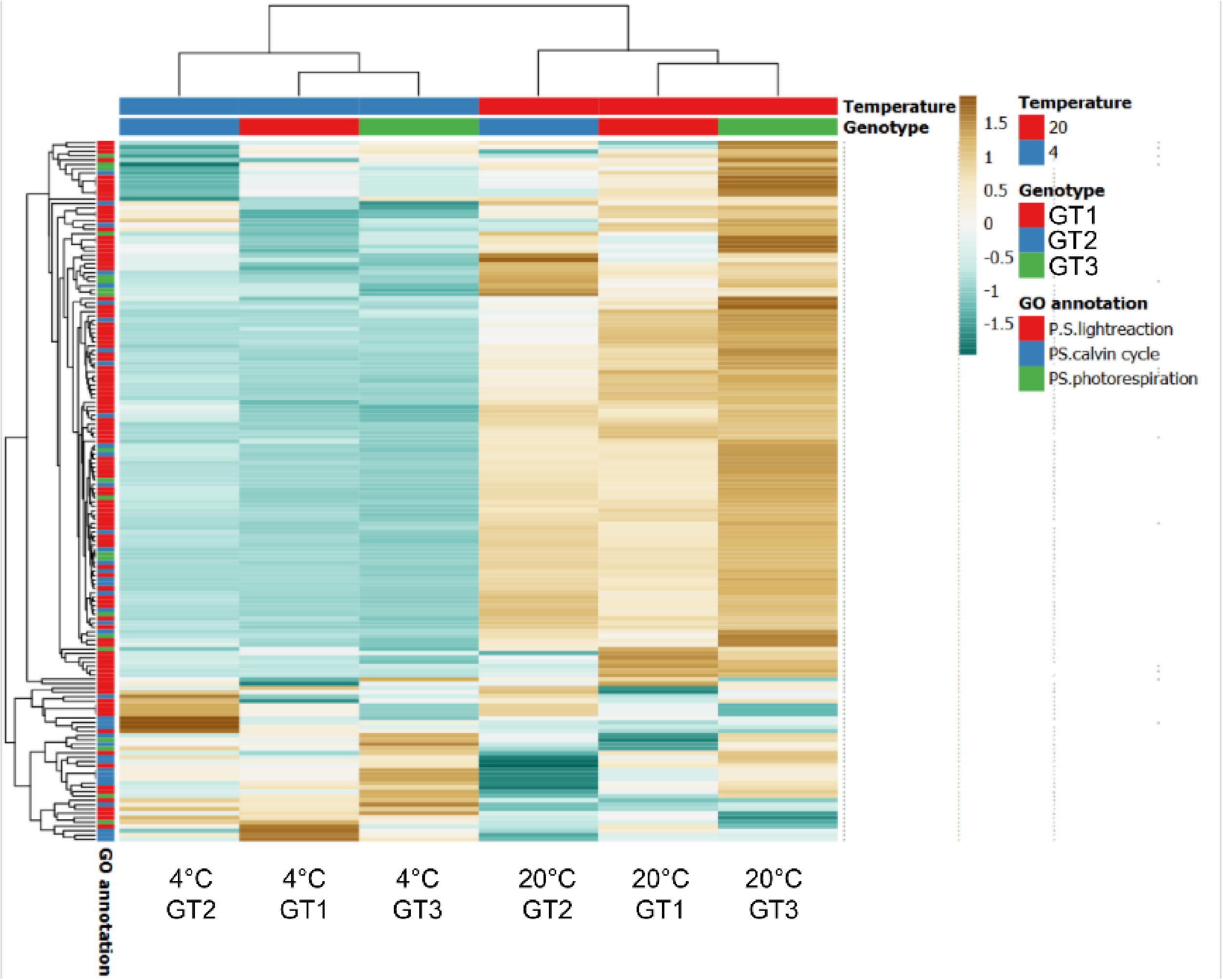


**Supplemental Figure 1.**
Heatmap analysis of 162 photosynthesis-related genes.
Unit variance scaling was applied and rows were clustered using Manhattan distance and average linkage. Columns are clustered using correlation distance and average linkage. This Figure supports Figure 2 from the main text.

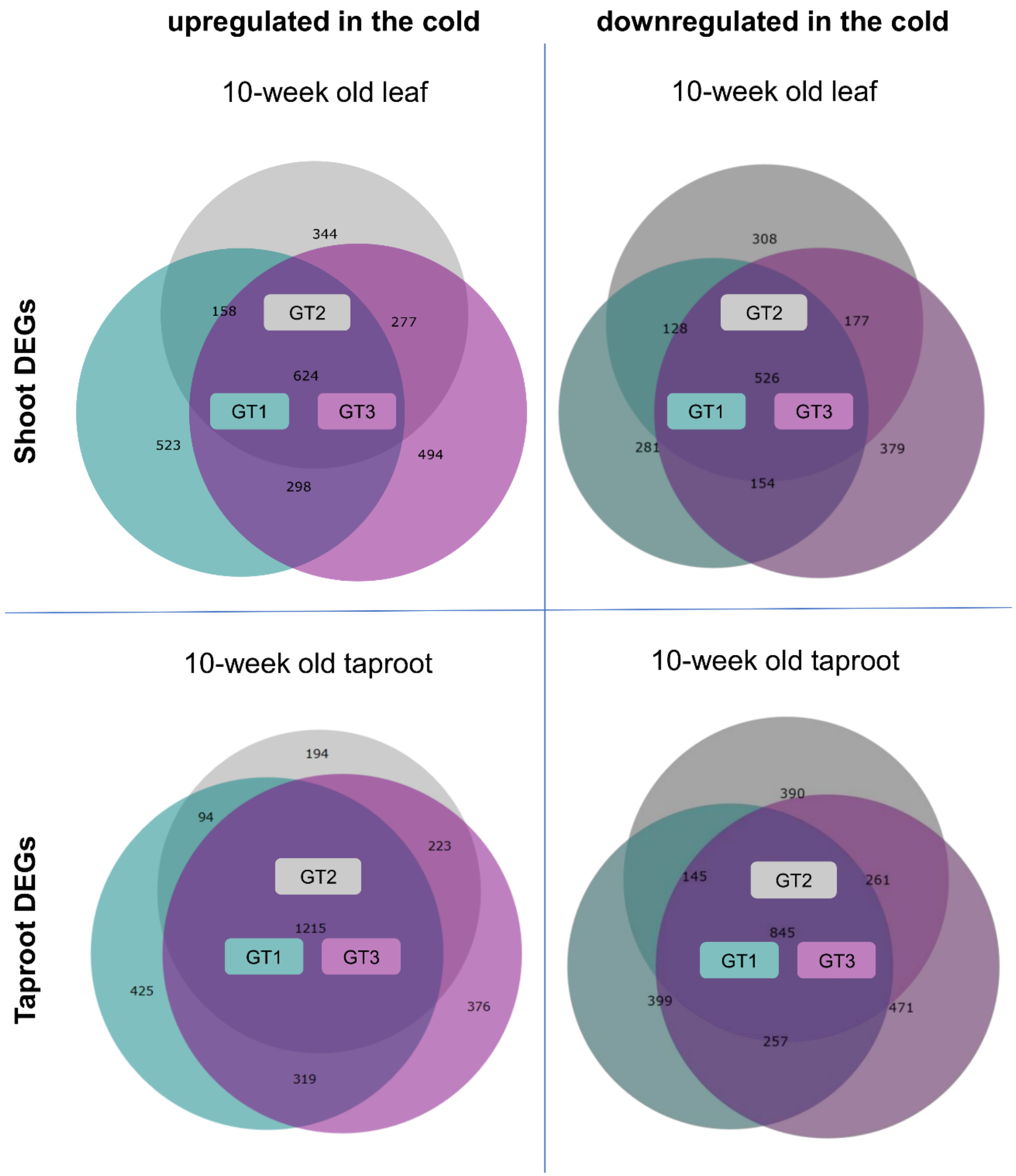


**Supplemental Figure 2.**
Venn diagrams of differentially expressed genes (DEGs) in leaves and taproots.
Numbers of up- (Log2 fold change ≥ 1) or down- (Log2 fold change ≤ -1) regulated genes (with a FDR ≤ 0.01) are given inside circles of Venn diagrams. The total number of common DEGs (i.e. in intersections of all genotypes) was higher in taproots than in shoots (1215 up- and 845 downregulated DEGs in taproots versus 624 up- and 524 downregulated in shoots). This figure supports Figure 2 from the main text.

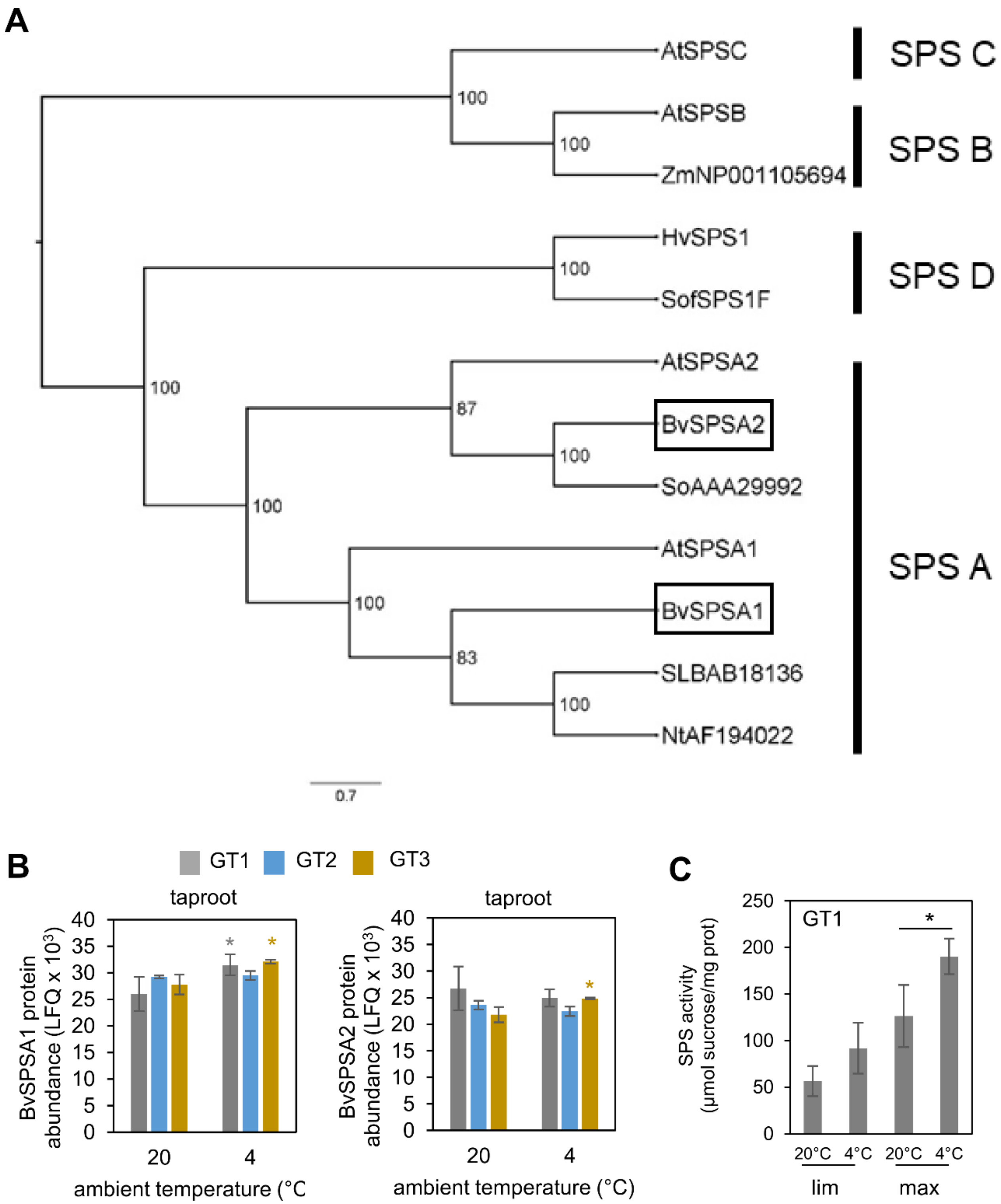


**Supplemental Figure 3.**
Phylogeny of Beta vulgaris SPS isoforms and protein abundance of BvSPS isoforms in taproots.
**(A)** Phylogeny of BvSPS proteins. **(B)** SPSA1 and SPSA2 protein abundance based on MS counts (label free intensities, LFQ units) from GT1, GT2, GT3 (BvSPSA1 = Bv2_030670_mgoq.t1; BvSPSA2 = Bv8_193450_doak.t1). **(C)** SPS activity in roots under substrate (F-6-P) limiting (lim) and maximum (max) conditions. This figure supports Figure 3 from the main text.

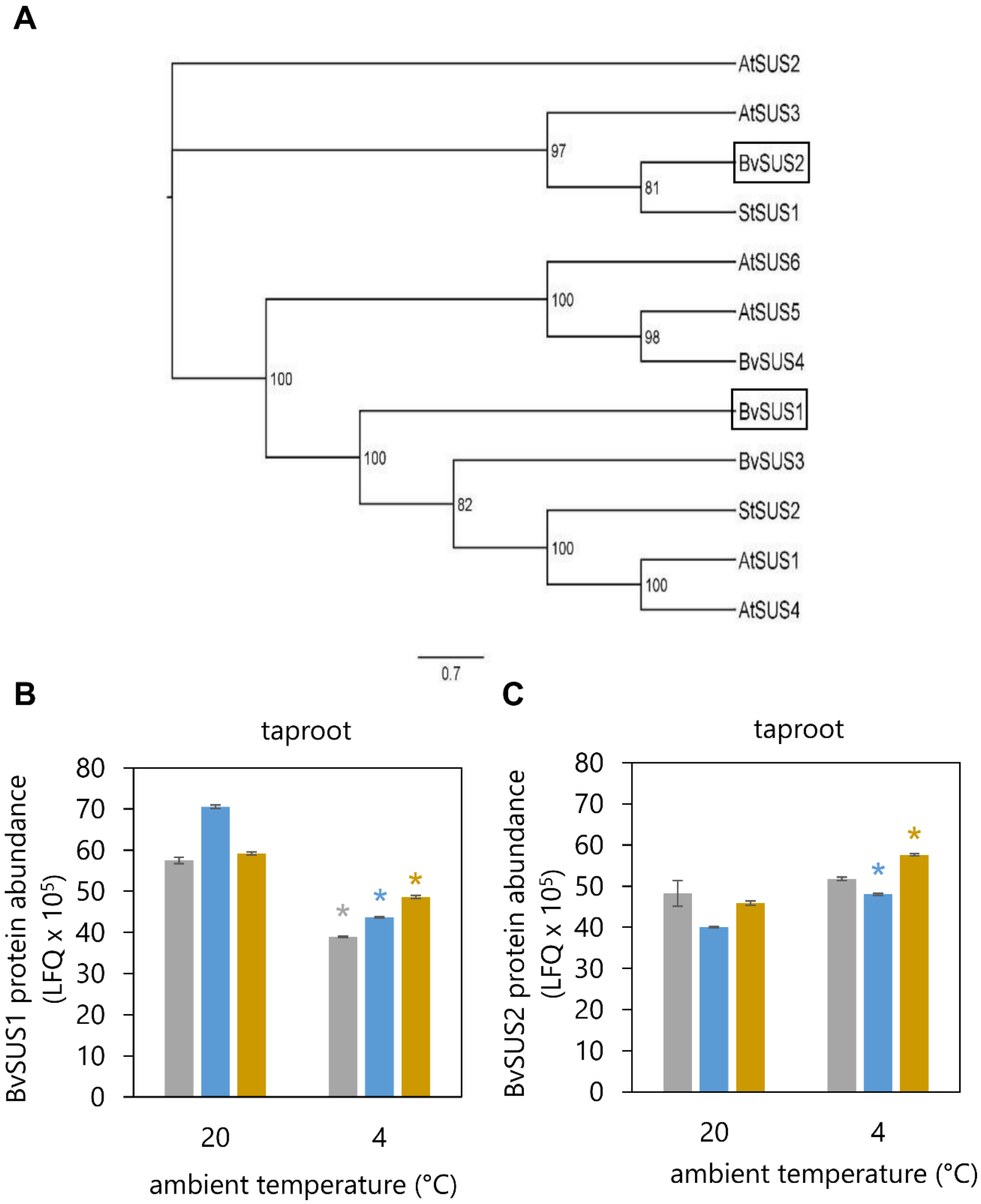


**Supplemental Figure 4.**
Phylogeny of Beta vulgaris SUS isoforms and protein abundance of BvSUS isoforms in taproots.
**(A)** Phylogenetic tree of sucrose synthase amino acid sequences from sugar beet, Arabidopsis and potato. Sugar beet proteins had the following identifiers: BvSUS1: Bv8_190960_nnjy.t1, BvSUS2: Bv7_163460_jmqz.t1, BvSUS3: Bv7_173620_ffuo.t1, BvSUS4: Bv4_084720_myet.t1. Arabidopsis proteins had the following identifiers: AtSUS1: AT5G20830, AtSUS2: AT5G49190, AtSUS3: AT4g02280, AtSUS4: AT3G43190, AtSUS5: AT5G37180, AtSUS6: AT1G73370. Potato proteins had the following identifiers: StSUS1: NP_001275237.1, StSUS2: XP_015166930.1. **(B, C)** BvSUS1 and BvSUS2 protein abundance (label free intensity) in soluble protein fraction of taproots from three different genotypes (white = GT1, blue = GT2, grey =GT3) grown at 20°C or grown at 20°C and transferred for two weeks to 4°C. This figure supports Figure 3 from the main text.

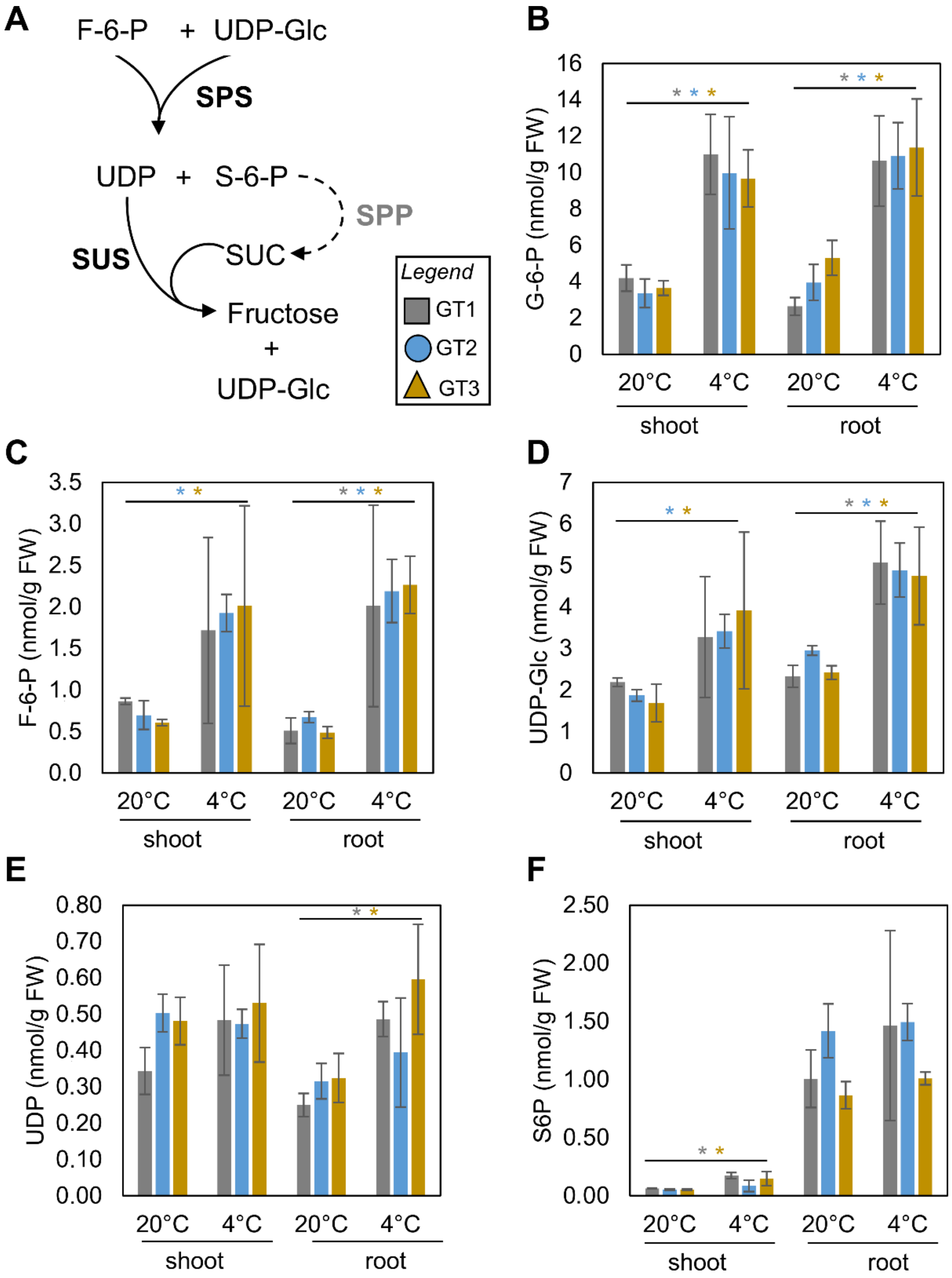


**Supplemental Figure 5.**

Phosphorylated metabolites in shoots and taproots of sugar beet plants.

**(A)** Schematic depiction of sucrose metabolizing processes. **(B-F)** Concentrations of phosphorylated metabolites in shoots and roots of three different genotypes (grey: GT1, blue: GT2, light brown: GT3) grown for 8 weeks under 20°C and then either kept for 2 more weeks at 20°C or transferred to 4°C. Abbreviations: SPS: Sucrose Phosphate Synthase, SPP: Sucrose Phosphate Phosphatase, SUS: Sucrose Synthase, G-6-P: Glucose-6-Phosphate, F-6-P: Fructose-6-Phosphate, UDP-Glc: UDP-Glucose, S-6-P: Sucrose-6-Phosphate. Data are means ± SD. Asterisks represent p-values < 0.05 according to double sided t-test in comparison to the values at control condition (20°C). This figure supports Figure 3 from the main text.

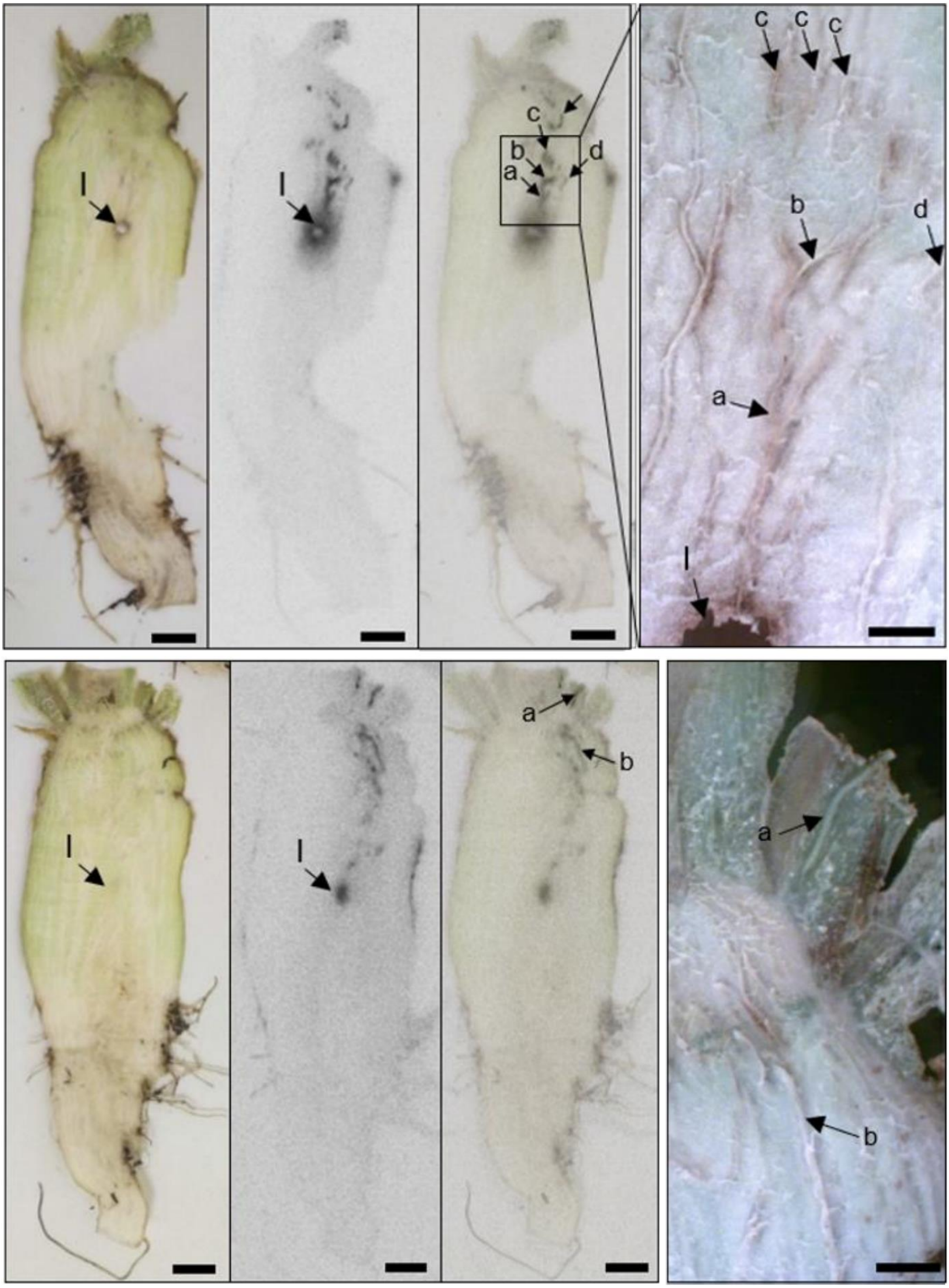


**Supplemental Figure 6.**

Exemplary pictures of radioactivity incorporated and distributed in taproot tissue in the cold.

Plants were grown for 10 weeks at 20°C and then transferred for 1 week to 12°C and for 1 week to 4°C. Taproots were inoculated with $^{14}C$-sucrose and harvested 5 days later. Thin longitudinal taproot slices were prepared by hand, pressed and dried. From left to right: photographic image, phosphor-imaging recording, overlay of photography and phosphor-image recording, magnification of section region of interest. Arrowheads point towards sites of radioactivity. Bars are = 5 mm for whole root pictures and 0.5 mm for magnifications (rightmost panels). This figure supports Figure 4 from the main text.

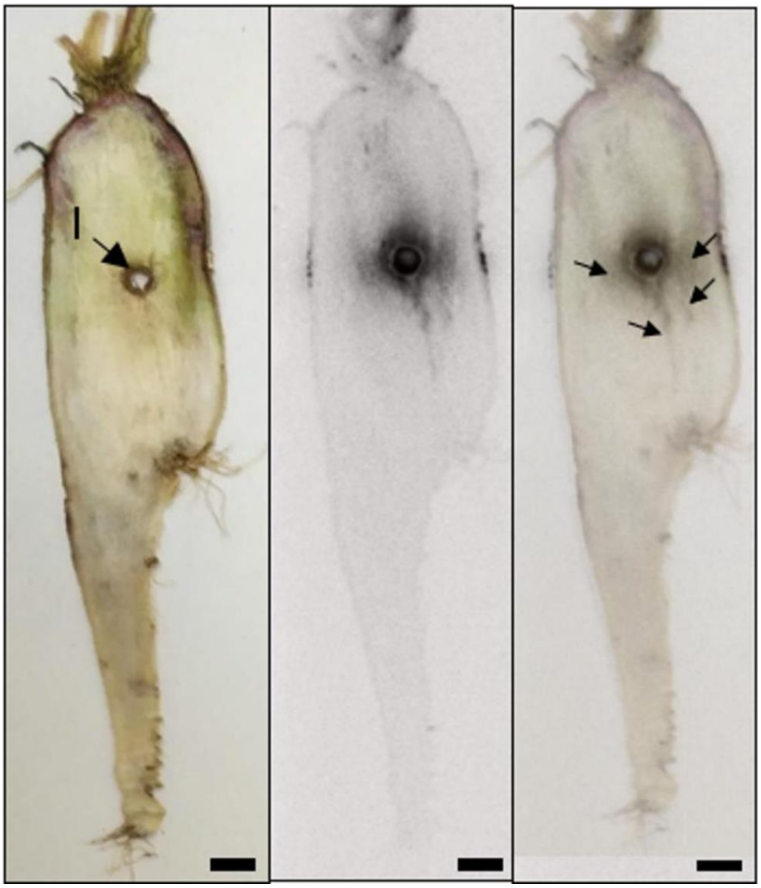


**Supplemental Figure 7.**

Exemplary pictures of radioactivity incorporated and distributed in taproot tissue.

Plants were grown for 10 weeks at 20°C and then taproots inoculated with $^{14}C$-sucrose and harvested 5 days later. Thin longitudinal taproot slices were prepared by hand, pressed and dried. From left to right: photographic image, phosphor-imaging recording, overlay of photography and phosphor-image recording. Arrowheads point towards sites of radioactivity. Bars are 5 mm. This figure supports Figure 4 from the main text.

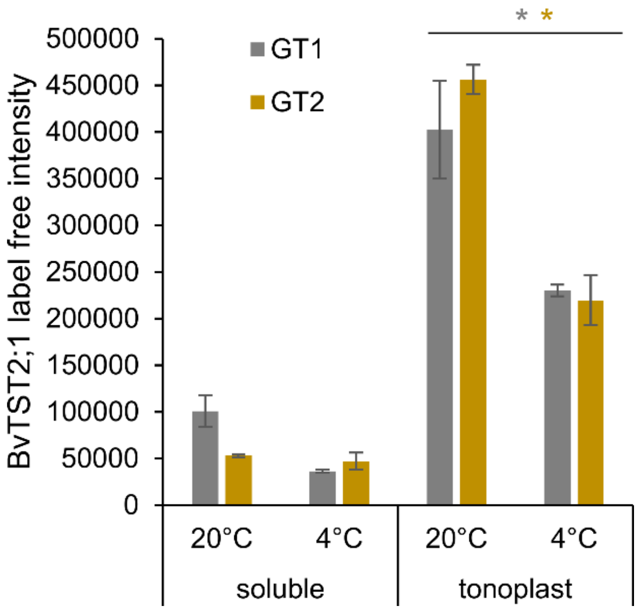


**Supplemental Figure 8.** Abundancy of BvTST2;1 protein levels. Protein abundance based on MS counts given as LFQ (label free intensity). Values represent means from n=6 biological replicates per genotype ± SE. Asterisks indicate significant differences between the 20°C and 4°C treatments according to t-test (* = $p < 0.05$). This figure supports Figure 5 from the main text.

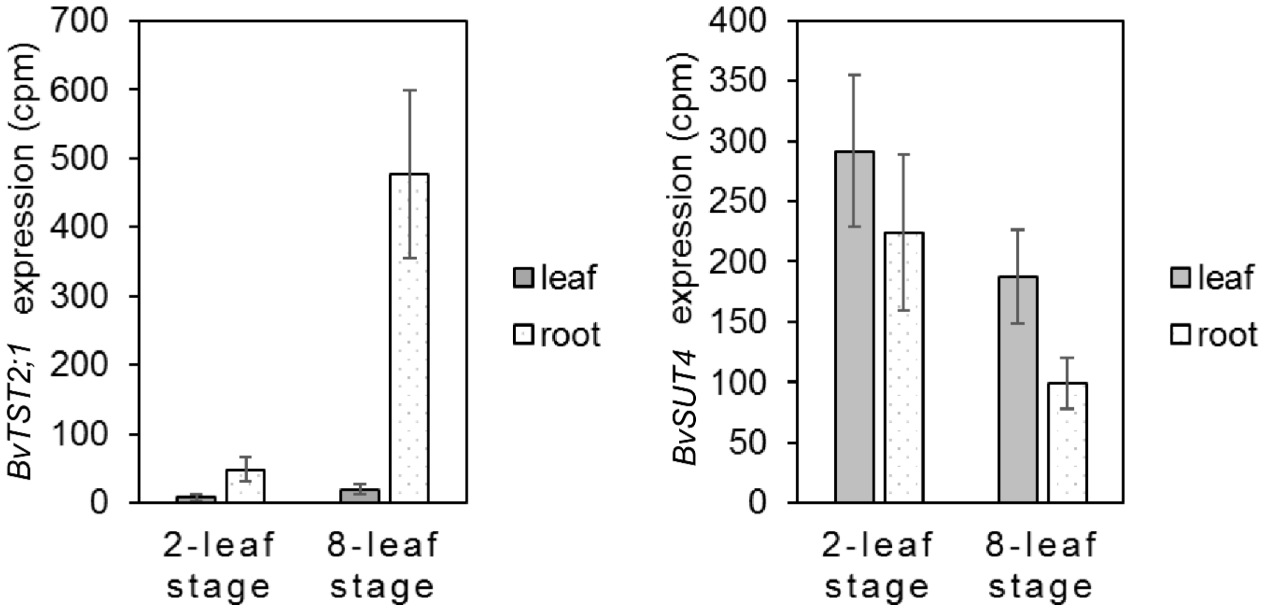


**Supplemental Figure 9.** Expression of *BvTST2;1* and *BvSUT4* in leaves and roots of sugar beet plants from the two and eight-leaf stage. This figure supports Figure 5 from the main text.

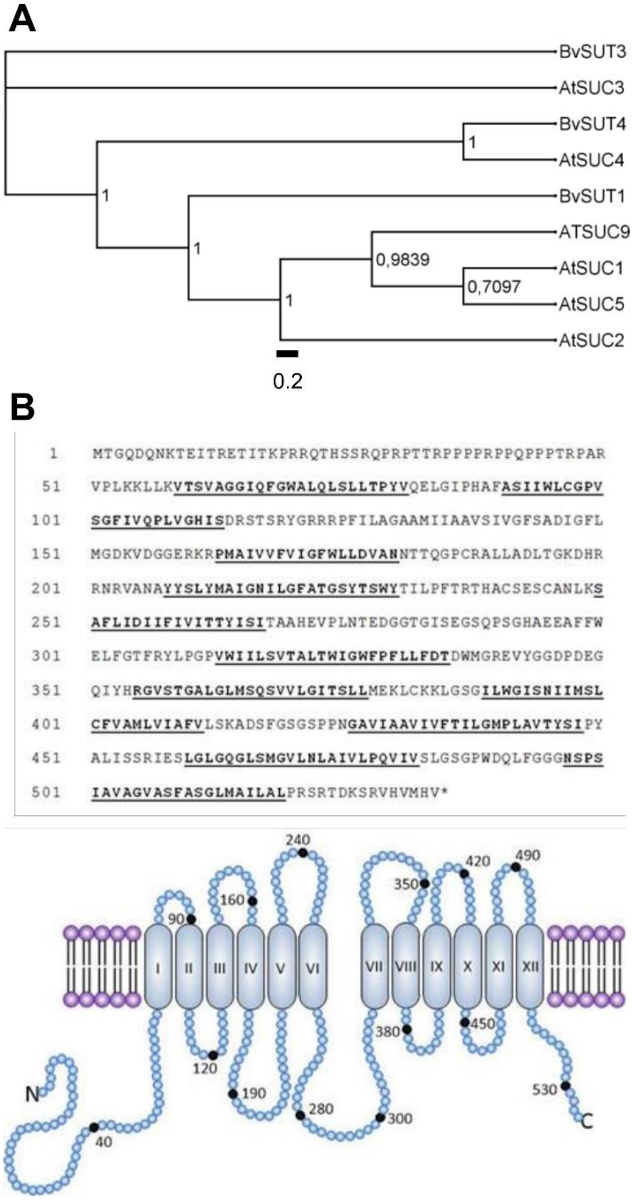


**Supplemental Figure 10.**
Phylogeny, sequence and predicted 2D-protein structure of BvSUT4.

**(A)** Unrooted phylogenetic tree of sucrose transporters from the SUT/SUC family of Beta vulgaris and Arabidopsis thaliana. Bayesian phylogenetic analysis was performed with MrBayes version 3.2.6 (Ronquist et al., 2012). MrBayes was run by conducting two parallel Metropolis coupled Monte Carlo Markov chain analyses four twenty thousand generations. The standard deviation of split frequencies was below 0.01. The tree was visualized using FigTree v.1.4.3. Sugar beet protein sequences had the following identifiers (RefBeet 1.2):
BvSUT1: Bv1_000710_gzum.t1,
BvSUT3: Bv6_154300_yemu.t1,
BvSUT4: Bv5_124860_zpft.t1.
Arabidopsis proteins had the following identifiers:
AtSUC1: AT1G71880,
AtSUC2: AT1G22710,
AtSUC3: AT2G02860,
AtSUC4: AT1G09960,
AtSUC5: AT1G71890,
ATSUC9: AT5G06170.

**(B)** Sequence and schematic depiction of the BvSUT4 protein. The protein has 535 aa and 12 transmembrane domains (underlined). The N-terminus includes the first 58 aa and the C-terminus the very last 14 aa, located in the cytoplasm of the cell. It has a central loop between transmembrane domain six and seven that includes 35 aa. This figure supports Figure 5 from the main text.
